# Single-cell transcriptomics identifies regulation of invasive behavior in *Drosophila* follicle cells with polarity loss

**DOI:** 10.1101/2022.06.09.495554

**Authors:** Deeptiman Chatterjee, Xian-Feng Wang, Allison Jevitt, Caique Almeida Machado Costa, Yi- Chun Huang, Wu-Min Deng

## Abstract

Apicobasal cell-polarity loss is a founding event in Epithelial-Mesenchymal Transition (EMT) and epithelial tumorigenesis, yet how pathological polarity loss induces plasticity changes remains largely unknown. To understand the mechanisms and mediators regulating plasticity upon polarity loss, we performed single-cell (sc) RNA sequencing of *Drosophila* ovaries, where inducing polarity-gene *l(2)gl* knockdown (Lgl-KD) causes invasive delamination of the follicular epithelia. Integrating Lgl-KD with the corresponding *wild-type* sc-transcriptome, we discovered clusters specific to various discernible phenotype-specific cell types and further characterized the regulons active in those cells. A genetic requirement of Keap1-Nrf2 signaling in promoting multilayer formation of Lgl-KD cells was further identified. Elevated expression of Keap1 increased the volume of delaminated follicle cells that undergo enhanced collective invasion via cytoskeletal remodeling. Overall, our findings describe the comprehensive transcriptome of the follicle-cell tumor model at the single-cell resolution and identify a previously unappreciated link between stress signaling and cell plasticity in early tumorigenesis.

## INTRODUCTION

Apical-basal cell polarity acts as a major gatekeeper against tumorigenesis in epithelial tissues (Royer and Lu, 2011). While its dysregulation is commonly associated with tumorigenesis (Rudrapatna, Cagan and Das, 2012; Chatterjee and Deng, 2019), cell-polarity disruption is tightly controlled to also allow cellular movement during critical developmental processes such as gastrulation and wound healing (Barriere et al., 2015). This regulation of cell polarity enables the cells to undergo changes in cell plasticity, which is the ability of cells to change their phenotype in response to environmental factors without acquiring genetic mutations. Plasticity in these cells facilitate Epithelial-to-Mesenchymal Transition (EMT) causing the cells to lose apical-basal polarity and weaken cell-cell adhesion with the purpose of promoting their movement beyond the confined space of the tissue (Moreno-Bueno, Portillo and Cano, 2008; Plygawko, Kan and Campbell, 2020). For these reasons, EMT is also associated with increased invasive behavior and metastatic migration of cancer cells (Mani et al., 2008; Brabletz et al., 2018). Recently emerging consensus in the field of EMT research draws attention to the presence of a continuum of metastable cells undergoing partial-EMT (pEMT) in cancer tissues, instead of one stable state of either epithelial or mesenchymal identity (Grigore et al., 2016; Saxena, Jolly and Balamurugan, 2020). A significant gap exists in our understanding of how the different metastable-cell states are regulated and how pEMT relates to the overall behavior of the invasive tumor tissue.

Epithelial polarity is maintained by the mutually-antagonizing, spatially-restricted protein complexes at the apical and basolateral cytosolic domains of the cell which are frequently found disrupted in several cancers (Huang and Muthuswamy, 2010; Elsum et al., 2012; Parker et al., 2014). A causative link between polarity loss and tumorigenesis has since been established in the fruit-fly *Drosophila melanogaster*, where the loss of basolateral polarity proteins – such as Scribble (Scrib), Discs large (Dlg) and Lethal giant larvae (l(2)gl or simply, Lgl) – combined with oncogenic Notch or Ras signaling, causes malignant tumorigenesis in several tissues (Papagiannouli and Mechler, 2004; St Johnston and Ahringer, 2010; Papagiannouli and M., 2013; Enomoto, Carmen and Igaki, 2018; Chatterjee and Deng, 2019). While several advances have since been made in identifying genetic factors that regulate tumorigenesis in the wing-disc tumor model (Doggett et al., 2015; Atkins et al., 2016; Dillard, Reis and Rusten, 2021; Logeay et al., 2022), progress toward developing an integrated understanding of the mechanisms driving neoplastic remodeling of the tissue is limited. This limitation is due to: (1) the use of bulk-tissue based approaches that cannot detect the spatial heterogeneity inherent to the growing tumor as well as large-scale changes in cell types, and (2) rampant malignancy in the wing-disc derived tumor model that obscures the detection of individual-cell behavior. In a recent study, we drew attention to the phenotypic heterogeneity presented within the follicular epithelia of *Drosophila* ovaries where Lgl-knockdown (Lgl-KD) induced polarity loss causes multilayer formation (Jevitt et al., 2021). Even in presence of oncogenic Notch signaling, the multilayered follicle-cell mass exhibits sustained growth and survival but do not show long-distance malignancy when transplanted into the host’s abdomen (Jevitt et al., 2021). This lack of malignancy provides an exceptional opportunity to study the phenotypic behavior of polarity-deficient cells and to identify how it is regulated by their underlying transcriptome.

The normally developing *Drosophila* follicular epithelium has been instrumental in our understanding of cellular plasticity that facilitates developmental EMT and collective migration of the anterior follicle cells and border cells (Silver et al., 2001). In 2019, we published a comprehensive sc-transcriptomic atlas describing gene-expression signatures of individual *w^1118^* follicle cells, also including the abovementioned cell types (Jevitt et al., 2020). In this study, we have utilized this *w^1118^* follicle-cell dataset as a template to identify cells that deviate from the normally-developing follicle-cell lineage as a consequence of inducing the functional loss of Lgl. We have validated the transcriptomic signatures of these deviated cells to characterize the distinct phenotypes that manifest in the ovaries with Lgl-KD in their follicle cells. We focused specifically on a cluster containing heterogenous cells that exhibit stress-associated regulon activity. A previously undiscovered role of the cytoprotective Keap1-Nrf2 signaling in regulating invasiveness of the follicle cells with polarity loss was identified and this invasive behavior was further characterized by staining for antibodies against relevant cytoskeletal proteins. Our findings in this study describe early gene-expression changes within dysplastic cells of the follicular epithelia and identifies novel signaling pathways that regulate the plasticity of delaminating tumor cells.

## RESULTS

### Lgl silencing causes distinct phenotypic and transcriptomic changes in egg chambers

Using the pan follicle-cell *traffic jam* (*tj*)-Gal4 driver with temperature-sensitive (TS) Gal80 (Gal80^TS^) repressor to control transcriptional induction, follicle-cell specific silencing of Lgl (*tj^TS^>lgl^RNAi^*) was induced in adult female flies for 72 hours (h), which resulted in the premature failure of oogenesis. This failure of egg development resulted in the accumulation of degenerated egg chambers within the epithelial sheath (Fig.1A). Additionally, phospho-Histone 3 (pH3) accumulation was observed in the follicle cells at the egg-chamber termini during mid-oogenesis, when control and lateral follicle cells cease to undergo mitosis (Fig.1A). Cross-section of these egg chambers revealed significant multilayering of the epithelia, where cells exhibited decreased expression of the differentiated follicle-cell marker Hindsight (Hnt) (Sun and Deng, 2007) and increased expression of the immature-cell marker Eyes absent (Eya) (López-Schier and St. Johnston, 2001; Sun and Deng, 2005), suggesting changes to the expected cell fate (Supplementary Fig.1). To determine how Lgl loss-of-function (Lgl-LOF) affects epithelial-cell plasticity, we stained them with common epithelial-cell markers Shotgun or Shg (also known as DE-Cad, the *Drosophila* homolog of E-Cadherin) and its binding partner Armadillo or Arm (α- Catenin homolog). Both Shg and Arm staining displayed gradually decreasing enrichment at the multilayered-cell junctions along the apical-basal axis, with the basal-most layer expressing Shg at levels comparable to that observed in the monolayered cells of the control egg chambers (Fig.1B). When the intensities of F-actin (Phalloidin) and Shg were measured along the apical-basal axis of the invasive multilayer shown in Fig.1B, significant decrease in Shg enrichment was observed along with mildly elevated F-actin at the apical-most delaminated cell (Fig.1C).

**Figure 1:**
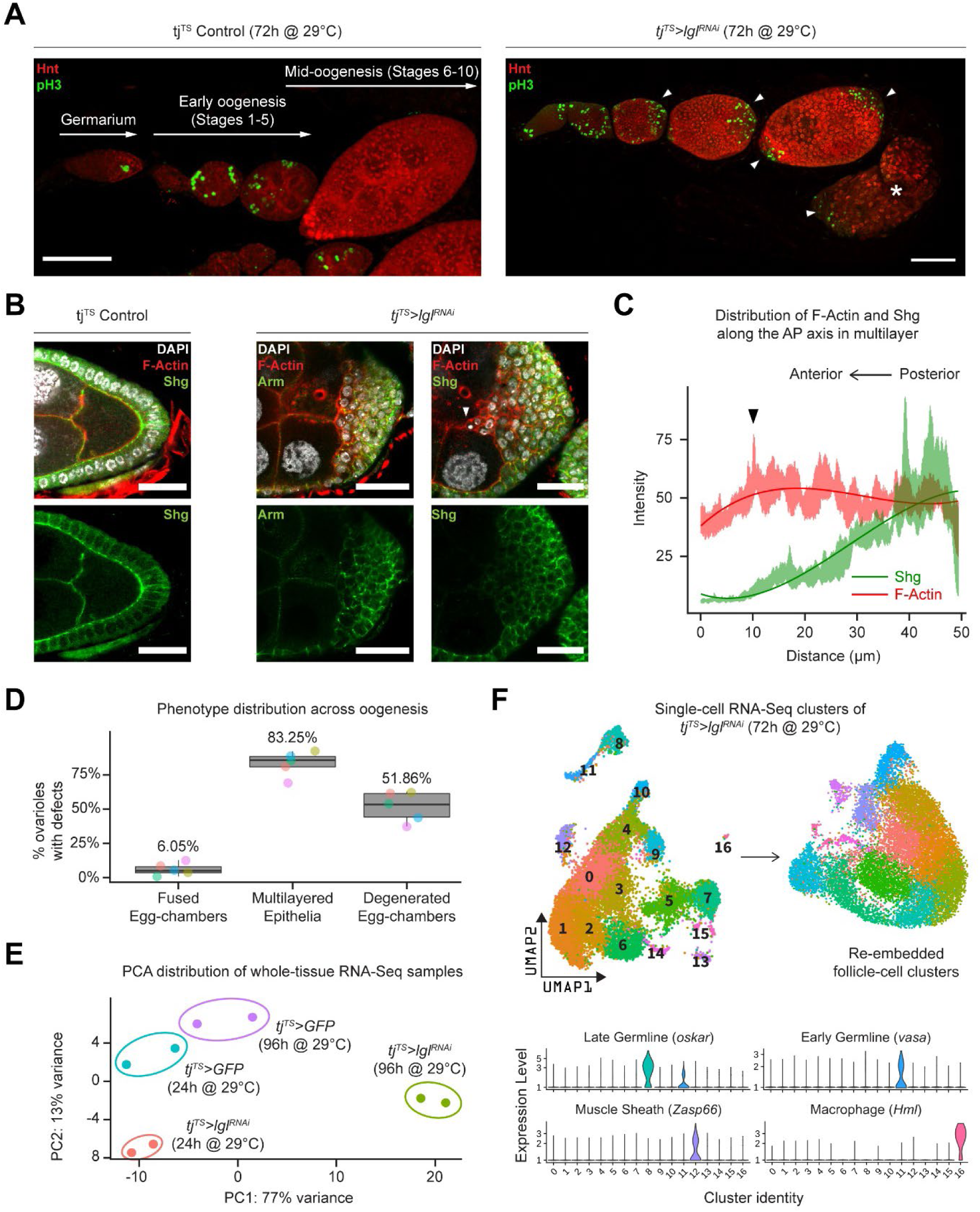
Inducing Lgl knock-down in follicle cells causes distinct phenotypic changes reflected in its transcriptome. **A.** (Left) Orthogonal projection of a single ovariole displaying individual egg chambers containing control follicle cells at early and midoogenesis. Follicle cells at mitotic stages are infrequently detected by pH3 staining (green), while endocycling follicle cells at midoogenesis are labeled by Hnt staining (red). (Right) Ovariole containing egg chambers with Lgl-KD in follicle cells exhibit continued cell division (marked by pH3 staining in green) in cells that accumulate at egg-chamber termini (arrowheads) at early-to-midoogenesis developmental transition and at midoogenesis. Degenerated egg chambers containing dying germline cells are marked by asterisks (*). Scale bars: 50 µm. **B.** (Left) Cross-section of the posterior egg-chamber epithelia containing control follicle cells that exhibit intact Shg (DE-Cad) staining at cell junctions. (Middle and Right) Posterior multilayers of egg chambers containing Lgl-KD in follicle cells show declining enrichment (green) of Armadillo (Arm; middle panel) and Shg (DE-Cad; right panel) along the anterior-posterior (AP) axis (right to left). F-Actin (red) is found enriched in cells at the apical-most layers; leading-edge of the invasive front is indicated by an arrowhead. Nucleus is marked by DAPI (white). Scale bars: 20 µm. **C.** Relative enrichment of F-Actin (red) and Shg (DE-Cad; green) along the AP axis across the biggest distance between the apical-most and basal-most cells in the multilayer shown in Fig.1B (right panel). Intensities are measured across a 10 µm thickness (Z-axis) and a trendline (Gaussian fit) is shown. The black arrowhead marks the leading-edge corresponding to that shown in Fig.1B. **D.** Box-and-whisker plot showing quantification of the different *tj^TS^>lgl^RNAi^* (72h in permissive temperature) phenotypes. Data was collected from 5 replicate trials (color-coded individually), consisting of 1,250 intact ovarioles from a total of 165 flies. **E.** PCA Plot showing the distribution of whole-tissue RNA-Seq samples for *tj^TS^* controls kept for 24h and 96h in permissive temperature and their experimental counterparts containing *tj^TS^>lgl^RNAi^*. Each uniquely-colored sample has two replicates that are grouped. **F.** Overview of the single-cell (sc) RNA-Seq workflow to isolate follicle-cell specific clusters from 14,537 *tj^TS^>lgl^RNAi^* ovarian cells, embedded on lower UMAP dimensions (top). Clusters containing nonepithelial cell-types are identified by the enrichment of specific markers (bottom).

Apically invasive movement was also detected in the single-cell derived *lgl^4^* MARCM clones, as well as groups of mitotic clones derived from homozygous *lgl^4^* LOF mutant follicle cells (Supplementary Fig.1). Quantifying our observations across 5 independent experiments, we found that multilayering at midoogenesis was the most prevalent (83.25%; n=280) large-scale phenotype observed across the different stages of oogenesis in *tj^TS^>lgl^RNAi^* ovarioles after 72h of Lgl-knockdown (72h-Lgl-KD) induction (Fig.1D). Additionally, about 6.05% (n=280) of egg chambers displayed fused egg chambers at early oogenesis, while degenerated egg chambers were observed in about half of all egg chambers (51.86%; n=280) at developmental stages beyond stage 9 or 10 during late oogenesis as a consequence of germline cell death (Fig.1D). Overall, these quantifications described the diverse egg-chamber phenotypes that manifest upon Lgl-knockdown (Lgl-KD) in follicle cells.

We next wondered if the diverse Lgl-KD phenotypes could also induce significant gene-expression changes in the ovaries. To test this, we performed whole-tissue RNA Sequencing (RNA-Seq) analysis of ovarian tissues from both control and Lgl-KD flies with RNAi induction for 24h (shorter) and 96h (longer periods of time). While 24h-Lgl-KD did not induce observable phenotypic changes, 96h-Lgl-KD displayed significant epithelial multilayering and germline-cell death (Supplementary Fig.2). Principal Component Analysis (PCA) comparing all four samples separated the 96h-Lgl-KD sample from others across PC1 (77% variance) suggesting significant difference between the sample with the differential phenotype vs the rest (Fig.1E).

When compared with all these “no-phenotype” samples collectively, several genes associated with late-stage oogenesis (e.g., *Vm26Ac*, *Femcoat*, *Cp15*, *yellow-g* etc.) showed reduced expression, while expression of genes involved in actin binding (Molecular Function GO:0003779; *p*=2.843x10^-3^), locomotion (Biological Process GO:0040011; *p*=3.748x10^-11^) and cell periphery (Cellular Component GO:0071944; *p*=2.904x10^-19^) were found elevated in the 96h-Lgl-KD sample (Supplementary Fig.2 and Supplementary Data 1). Additionally, elevated expression of apoptotic-cell clearance genes, such as *croquemort* (*crq*) (0.66 log_2_FC; *p*=0.00015) and *draper* (*drpr*) (0.642 log_2_FC; *p*=0.00027) was also detected in the 96h-Lgl-KD samples, possibly in response to the germline-cell death (Franc et al., 1996, 1999; Etchegaray et al., 2012). While these results mostly support the observed phenotypic changes, they do not distinguish gene expression by the distinct phenotypes or individual cell types. Therefore, to identify transcriptional changes specific to follicle cells, single-cell RNA-Seq (scRNA-Seq) was performed on cells isolated from 72h-*tj^TS^>lgl^RNAi^* ovaries.

### scRNA-Seq analysis identifies unique clusters associated with Lgl-KD dataset

On the basis of gene-expression similarities, 16,060 ovarian cells of the *tj^TS^>lgl^RNAi^* dataset were grouped into 17 clusters (Fig.1F and Supplementary Fig.3). Follicle cells were isolated for more targeted analyses by positively identifying non-epithelial cell types using previously described markers (Jevitt et al., 2020) (*vasa*+ and *oskar*+ early and late germline cells, *Zasp66*+ muscle sheath cells and *Hml*+ macrophage-cell population known as the hemocytes) and removing them from the dataset. The remaining 14,537 Lgl-KD follicle cells were then integrated with 17,875 *w^1118^* follicle cells (that were processed similarly to remove non-epithelial cells) and were collectively assembled into 20 clusters (Fig.2A). These clusters were annotated according to the differential enrichment of stage- and cell-type specific marker expression (Fig.2B). Consistent with the phenotypic observation that egg chambers degenerate post-midoogenesis around stage 8, clusters of cells belonging to egg chambers at stage 9 and beyond (clusters 9, 10, 11, 14 and 19) had lower proportions of Lgl-KD cells (Fig.2C-D). Significantly, we also identified clusters (clusters 7, 8, 13, 16 and 17) that were unique to the Lgl-KD dataset (Fig.2E). Our integration based approach was thus able to separate clusters unique to the individual datasets, possibly representative of cells exhibiting the diverse stage-specific Lgl-KD phenotypes and the loss of cells from expected stages comparable to the *w^1118^* developmental lineage.

**Figure 2:**
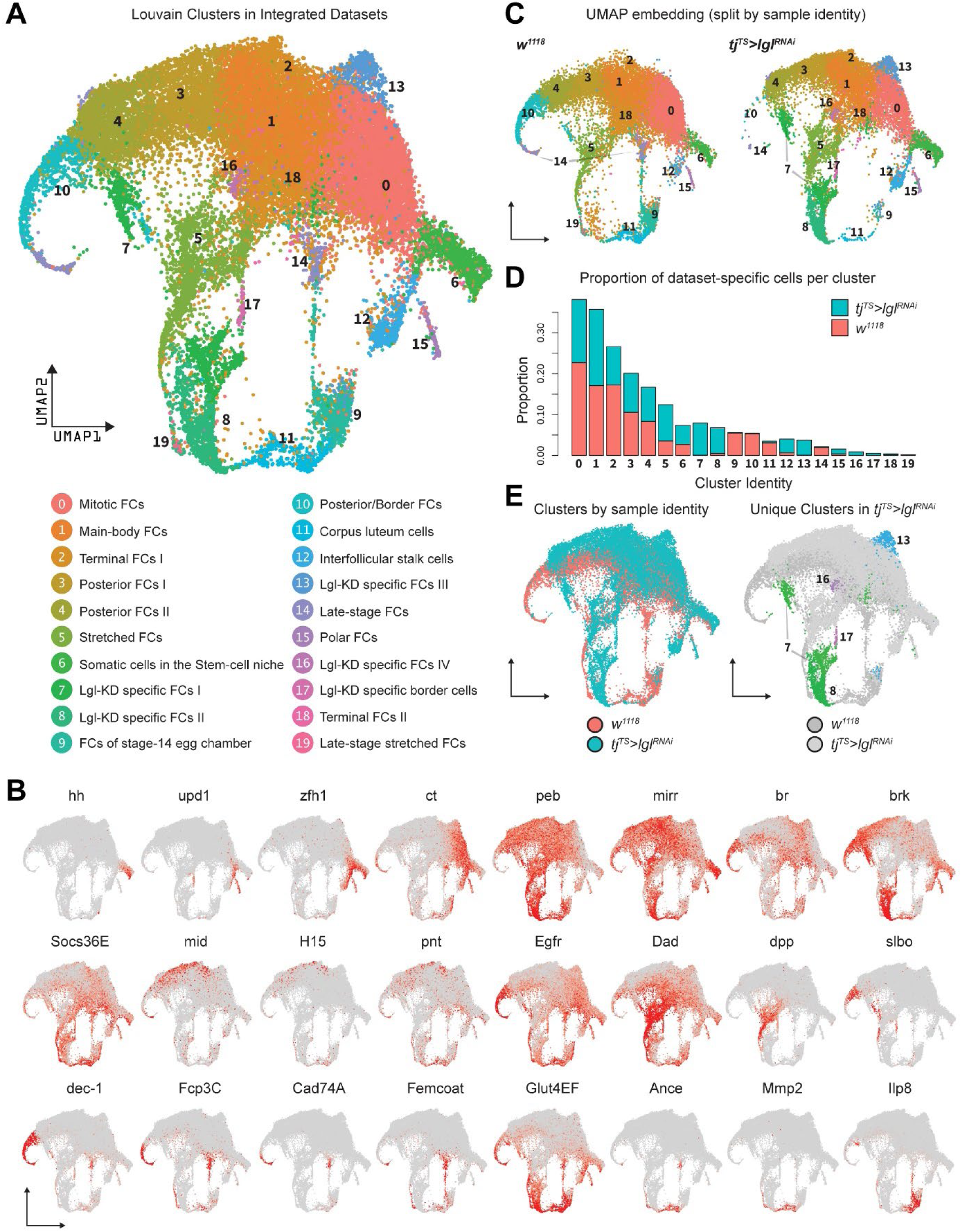
Integration of *w^1118^* and *tj^TS^>lgl^RNAi^* sc-datasets identifies clusters unique to the *tj^TS^>lgl^RNAi^* follicle cells. **A.** UMAP plot of 17,874 *w^1118^* follicle cells integrated with 12,923 *tj^TS^>lgl^RNAi^* follicle cells, grouped into 20 clusters, with their approximated identities listed below. **B.** Canonical marker expression used to annotate the 20 integrated clusters reveal cluster-specific identities of the somatic cells of the stem-cell niche (*hh*), polar follicle cells (*upd1*), stalk cells (*zfh1*), immature mitotic follicle cells (*ct*), mature post-mitotic cells (*peb*), main-body follicle cells (*mirr* and *br*), terminal follicle cells (*brk*, *Socs36E* and *Egfr*), posterior follicle cells (*mid*, *H15* and *pnt*), stretched cells (*Dad* and *dpp*), border cells (*slbo*), follicle cells of the vitellogenic (*dec-1*) and choriogenic stages (*Fcp3C*, *Cad74A*, *Femcoat* and *Glut4EF*), and the cells of the terminally- fated Corpus luteum (*Ance*, *Mmp2* and *Ilp8*). **C.** Distribution of cluster-specific cells in the UMAP embedding is shown split by their dataset of origin. **D.** Bar plot showing the proportion of cells in each cluster. Each bar is further divided by the dataset of origin (*w^1118^* is represented by salmon color and *tj^TS^>lgl^RNAi^* by teal). **E.** UMAP plot showing the overlap between cells belonging to each dataset (left) and unique clusters in the *tj^TS^>lgl^RNAi^* dataset (right).

Keeping cluster identities intact, the *w^1118^* cells were removed and the 14,537 Lgl- KD follicle cells were re-embedded in the same UMAP space that was built using anchor-restricted principal components upon integration (Fig.3A, left). The underlying lineage transitions among the cells of unique Lgl-KD clusters were then estimated by dynamically modeling their inherent RNA velocity (Bergen et al., 2020). The resultant stream of velocity vectors placed clusters 7, 13, 16 and 17 along a linear transition that terminated into cluster 8, while cluster 8 itself showed mixed lineage as was suggested by the non-uniform direction of velocity vectors (Fig.3A, right). The latent time experienced by the cells in these clusters as well as the likely terminal end-points were then inferred from the estimated lineage, which subsequently suggested that clusters 13, 16 and the near-terminal cells of cluster 7 were the stable end- points of the assumed lineage while cluster 7 displayed highly dynamic transcriptional transition (Fig.3B). Overall, given the relaxed topology of certain cluster manifolds as well as the results from our linage analysis, we concluded that the unique clusters were a mix of stable terminal cell states as well as transitioning cell states.

**Figure 3:**
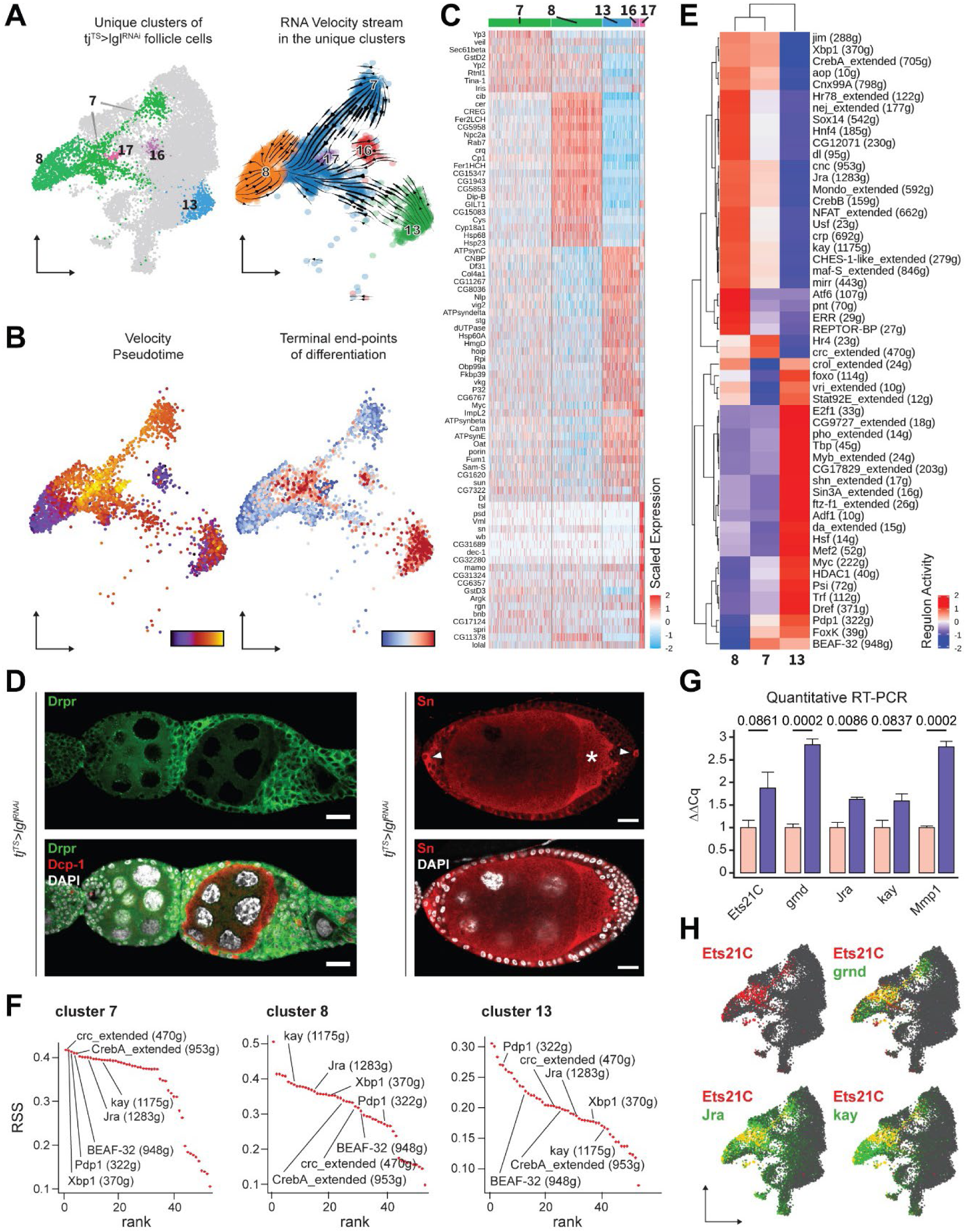
Unique clusters of *tj^TS^>lgl^RNAi^* dataset exhibit distinct gene-expression and regulon activity. **A.** (Left) UMAP plot of re-embedded *tj^TS^>lgl^RNAi^* follicle cells with the unique clusters being highlighted. (Right) Stream of RNA velocity vectors are superimposed on the unique *tj^TS^>lgl^RNAi^* clusters revealing the underlying lineage. **B.** (Left) Cells are colored according to their arrangement on the inferred pseudotime from early (purple) to late (yellow). (Right) Probability of cells at terminal states of the inferred lineage, where stable transcriptional end-points are colored in red and root cells are colored in blue. **C.** Heatmap of the top 20 specifically expressed markers (if present) in each unique cluster of the *tj^TS^>lgl^RNAi^* dataset. Selected genes are expressed in a minimum of 75% cells per cluster. Range of gene expression is scaled within +2 (red) to -2 (blue) log_2_ fold change. **D.** Confocal images showing Drpr (Left) in green and Sn staining (Right) in red. Dcp-1 (red) is also shown in left panel to mark dying germline cells within degenerating egg chambers. Nucleus is marked by DAPI (white). Scale bars: 20 µm. **E.** Heatmap of the scaled (and centered) activity scores of regulons in clusters 7, 8 and 13. Both columns (clusters) and rows (regulons) are clustered hierarchically. **F.** Regulon Specificity Score (RSS) Plot showing the cluster-specific ranks of regulon enrichment for clusters 7, 8 and 13. **G.** Bar Plot showing relative expression of mRNAs of JNK signaling components using quantitative RT-PCR. Bars representing control is colored salmon while *tj^TS^>lgl^RNAi^* is colored purple. **H.** Gene enrichment plots for Ets21C (red). *Jra*, *kay* and *grnd* are colored green, while overlapping enrichment is colored yellow.

### Unique Lgl-KD clusters exhibit cluster-specific gene expression and regulon activity

To characterize clusters 7, 8, 13, 16 and 17 further, we identified their differentially expressed markers (Fig.3C). Apoptotic-cell clearance markers *drpr* (Etchegaray et al., 2012) and *crq* (Franc et al., 1999) were specifically enriched in cluster 8, while polar and border-cell associated markers (Jevitt et al., 2020), such as *unpaired* (*upd1*), *slow border cells* (*slbo*) and *singed* (*sn*), were found enriched in cluster 17. We validated the expression of *sn* and *drpr* within the multilayers and found them to be expressed in the polar follicle cells within the multilayer and in the follicle cells of egg chambers that associate with dying germline cells respectively (Fig.3D). Border-cell specific expression of *sn* further implied that once the egg chambers have individualized, the polar to border-cell fate is not disrupted upon Lgl-KD, despite the impairment of border-cell migration. In contrast, clusters 7, 13 and 16 did not exhibit distinguishable markers. While markers of immature, mitotic follicle cells (Jevitt et al., 2020) such as *Myc*, *Df31* and *HmgD* were detected in cluster 13, it showed significant overlap with cluster 16, with the notable exception of the *Drosophila* Cdc25 homolog *string* (*stg*) expression. The expression of *stg* being higher in cluster 13 than that in cluster 16, we concluded that the clusters 13 and 16 likely represented mature (endocycling) and immature (mitotic) follicle-cell states, respectively, of the same cell-type identity. Additionally, very few specific markers were identified for cluster 7, as the markers either shared gene expression with (1) the main-body follicle cells (expressing *Yp2* and *Yp3*), suggestive of a shared (and possibly, derivative) lineage, and (2) clusters 8, 13 and 16, exhibiting only quantitative differences between them. These observations highlight the limitations of marker validation to identify specific cells of the differential Lgl-KD phenotype.

We used SCENIC (Aibar, 2017) to further identify the active regulons in the clusters unique to the Lgl-KD dataset: cluster 7 (1,144 cells), cluster 8 (913 cells) and cluster 13 (538 cells). We removed clusters 16 and 17 from this analysis due to the significantly low cell numbers (128 and 79 cells, respectively) in those clusters. While clusters 7 and 13 showed major differences in the enriched regulon networks, we found only 2 regulons (Hr4 and crc_extended) specific to cluster 7 (Fig.3E). Hierarchically grouping the clusters based on scaled regulon activity found cluster 7 to be more similar to cluster 8 than cluster 13, suggesting significant overlap between the regulons active in their constituent cells. To further explore the relatedness between the clusters, we assessed the Regulon Specificity Score (RSS) (Suo et al., 2018) of each regulon for individual clusters (Fig.3F and Supplementary Data 3). The top 5 regulons of cluster 7 – crc_extended (RSS=0.42), Xbp1 (0.42), Pdp1 (0.41), BEAF-32 (0.41) and CrebA_extended (0.41) – were identified along with the regulons associated with AP-1 transcription factors *Jra* (0.4) and *kay* (0.39) that have been reported to drive tumorigenic JNK signaling upon polarity loss. Comparable scores for kay (0.39), Jra (0.37) and Xbp1 (0.36) and low scores for CrebA_extended (0.35), crc_extended (0.33), BEAF-32 (0.32) and Pdp1 (0.27) were observed for cluster 8, while cluster 13 showed lower scores overall for each regulon. Interestingly, cluster 13 and cluster 8 exhibited inverse ranking of these regulons with Pdp1 scoring the highest (0.27) and kay scoring the lowest (0.17) RSS scores in cluster 13, highlighting a dynamic transition of regulon activity along the inferred lineage. Collectively, SCENIC was able to detect the common as well as distinct transcriptomic states of the cells in unique Lgl-KD clusters, while also highlighting the heterogeneity among them.

Given the discovery of the JNK signaling associated AP-1 transcription factors, *Pdp1* and *ftz-f1*, that have been previously implicated in polarity-loss induced metastatic-tumor formation in the wing discs (Igaki, Pagliarini and Xu, 2006; Uhlirova and Bohmann, 2006; Bunker et al., 2015; Kulshammer et al., 2015), we sought to assess the presence of tumorigenic JNK signaling in the Lgl-KD follicle-cell model. Using quantitative Real-Time (qRT)-PCR, we specifically detected transcripts of the tumorigenic JNK signaling components (Uhlirova and Bohmann, 2006; Andersen et al., 2015; Toggweiler, Willecke and Basler, 2016) such as Ets21C, TNFα-receptor Grindelwald (grnd), Jra, kay and the downstream target Matrix metalloproteinase-1 (Mmp1) in whole ovaries with Lgl-KD in follicle cells. Compared to the control ovaries, we detected significant upregulation of *grnd* (GAPDH-normalized expression relative to control: 2.84 ± 0.122 Standard Error; *p*=0.0002), *Jra* (1.62 ± 0.059; *p*=0.0086) and *Mmp1* (2.79 ± 0.129; *p*=0.0002) and noticeable upregulation of *Ets21C* (1.88 ± 0.35; *p*=0.0861) and *kay* (1.58 ± 0.188; *p*=0.0837) in ovaries with Lgl-KD follicle cells (Fig.3G). When we mapped the expression of *Ets21C*, *grnd*, *Jra* and *kay* to the UMAP-embedded Lgl-KD follicle cells, we found that while both clusters 7 and 8 exhibit *Jra*, *kay* and *grnd* enrichment, *Ets21C* is specifically detected in cluster 7, the terminal states of which also overlapped with dynamic *Mmp1* expression (Fig.3H and Supplementary Fig. 4). Taken together, our results suggest that the highly-transient cluster 7 consists of tumorigenic cells and so we next focused on smaller neighborhoods within this cluster to identify markers that could help us detect the precise location of these cells.

### Cluster 7 represents transient cells with heterogenous gene expression

To explore cluster 7 heterogeneity, we subdivided its 1,144 cells even further and obtained 5 transcriptomically similar neighborhoods (Fig.4A). The underlying lineage among these subclusters was then inferred from the inherent RNA velocity specific to cells in cluster 7. From the stream of velocity vectors superimposed on the UMAP embedding, we deduced that clusters 7_0 (293 cells) and 7_2 (193 cells) were the starting points of the inferred lineage that terminated into cluster 7_1 (291 cells), while clusters 7_3 (191 cells) and 7_4 (176 cells) were the intermediate states. To systematically describe the regulatory relationships of transcription factors in all cluster 7 cells, we compared the similarity of regulon activity scores for all possible regulon pairs based on the Connection Specificity Index (CSI) matrix (Bass et al., 2013). The 16 regulons active in cluster 7 (including those that incorporate low confidence transcription-factor associations) were organized into at least 2 larger clusters (Fig.4B). One of these highly correlated clusters contained the AP-1 transcription-factor heterodimers *Jra* and *kay* along with the basic Helix-Loop-Helix (bHLH) transcription factor *Usf, ftz transcription factor 1* (*ftz-f1*) and the GATA transcription factor *serpent* (*srp*). The other cluster included the AP-1 interacting gene *Activating transcription factor 3* (*Atf3*), Nuclear Receptor transcription factors *Hormone receptor 4* (*Hr4*) and *Hepatocyte nuclear factor 4* (*Hnf4*), the bHLH transcriptional repressor *hairy* (*h*), *Cyclic-AMP response element binding protein A* (*CrebA*) and *Ets at 21C* (*Ets21C*). Interestingly, these regulons showed distinct cluster-specific activity, with significantly different activity scores for each regulon per cluster (Fig.4C). Taken together, our results from the regulon analysis in cluster 7 further concluded that the heterogenous cells of this cluster were regulated by hierarchically organized, overlapping regulatory modules that work together to characterize the underlying transitioning cell state.

**Figure 4:**
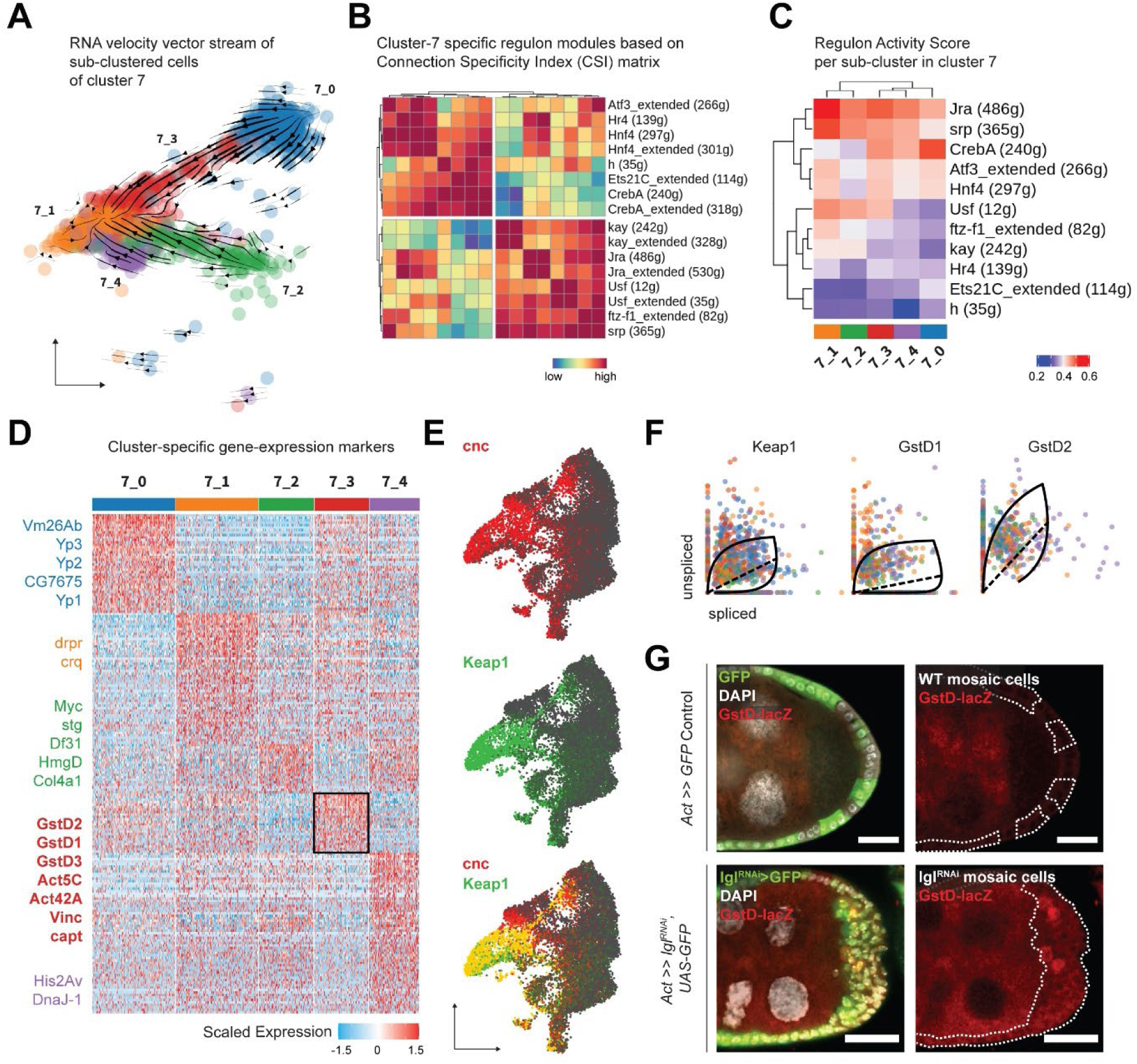
Cluster 7 cells exhibit hierarchically-organized regulon activity and gene- expression heterogeneity. **A.** UMAP Plot of subdivided cluster 7 cells, superimposed with RNA velocity vectors. **B.** Correlation heatmap representing the Connection Specificity Index (CSI) matrix of active regulon modules. **C.** Heatmap of the unscaled activity scores of regulons in the subclusters of cluster 7. **D.** Heatmap of the differentially expressed markers in each subclusters of cluster 7 cells, scaled within +1.5 (red) and -1.5 (blue) log_2_ fold change. **E.** Gene enrichment plot for *cnc* (red), *Keap1* (green) and their overlap (bottom). **F.** Phase portraits showing dynamic behavior of genes in cluster 7 cells (colored according to subcluster identity as shown in Fig.4A). Solid line represents the learned splicing dynamics while the dotted line represents the inferred steady state of gene expression. *Keap1*, *GstD1* and *GstD2* exhibits increased transcription in cluster 7. **G.** Reporter expression of *GstD-lacZ* is detected by β-Gal expression (red) within the multilayered cells of *lgl^RNAi^* follicle cells (green; clonal boundaries are marked by the dotted white lines). Nuclei is marked by DAPI (white). Scale bars: 20 µm.

Next, we identified the differentially enriched markers of the different subclusters of cluster 7 to further characterize its underlying heterogeneity. Differential gene expression analysis identified cluster 7_0 as mature, post-mitotic follicle cells (*Yp1-3*+ and *Vm26Ab*+) and cluster 7_2 as immature, mitotic follicle cells (*stg*+, *Myc*+ and *HmgD*+), while cluster 7_1 expressed markers involved in apoptotic-cell clearance (*drpr*+ and *crq*+) representing the degenerated egg chambers (Fig.4D). The inferred trajectory of cluster 7 therefore signified the dynamic transcription that the heterogenous population of Lgl-KD cells (defined by cell-fate differences) experienced overtime. We were particularly interested in identifying the differentially-expressed markers of cluster 7_3 which transitioned from cluster 7_0 representing mature follicle-cell type, as these genes would likely represent the invasive front of the multilayered cells that is composed of delaminating post-mitotic follicle cells. We found several relatively enriched markers of cluster 7_3 that associated with the actomyosin cytoskeleton (*Act5C*, *Act42A*, *Vinc*, *capt*, *cib* and *sqh* and *Mlc-c*), were part of the glutathione metabolic process (*GstD1*, *GstD2* and *GstD3*), Rab protein signal transduction (*Rab1*, *Rab7* and *RabX*), as well as the lysosomal trafficking pathway (*Rab7*, *cathD*, *Cp1*, *Cln3* and *Swim*) (Supplementary Data 4). Genes coding for the cytosolic Glutathione S-transferase (Gst) family proteins are expressed under the control of the transcription factor *cap-n-collar* or *cnc* (*Drosophila* Nrf2 homolog) (Sykiotis and Bohmann, 2008), that was found with high confidence in the *Jra*, *kay* and *Usf* associated regulons and with low confidence in the Atf3_extended regulon (Supplementary Data 5). From our results, we concluded that cluster 7_3 likely represented cells with elevated activation of the *cnc*-driven, stress-responsive Keap1-Nrf2 signaling pathway (Yamamoto, Kensler and Motohashi, 2018).

We detected an enrichment of *cnc* and its endogenous inhibitor *Keap1* in clusters 7 and 8 of the Lgl-KD dataset (Fig.4E). In support of the transient expression inferred from detecting the elevated expression of Keap1-Nrf2 pathway target genes in cluster 7_3, RNA velocity analysis detected the dynamic behavior of *Keap1* as well as several Gst genes (Fig.4F). Using the *GstD-lacZ* enhancer-trap reporter assay to validate Keap1-Nrf2 signaling activation (Sykiotis and Bohmann, 2008), we detected infrequent *β-Gal* activity in subsets of cells within the multilayer (Fig.4G). We were therefore able to conclude that cluster 7 likely represented the heterogenous cells of the multilayer, where the dynamic activation of Keap1-Nrf2 signaling was detected in a smaller subset of those cells.

### Keap1-Nrf2 stress signaling regulates invasive multilayering

To determine how Keap1-Nrf2 signaling affects epithelial multilayering, we knocked down the expression of its upstream components *cnc* and *Keap1* in Lgl-KD cells (Fig.5A-B). In ovaries with Lgl-KD follicle cells, 86.35% (N=793) ovarioles contained egg chambers with more than two layers of cells at mid-oogenesis, whereas Lgl-KD+cnc-KD ovaries exhibited comparable multilayering in only 27.2% ovarioles (N=964). Keap1-KD in Lgl-KD follicle cells exhibited a partial rescue of the Lgl-KD phenotype as only 35.46% ovarioles (N=897) contained egg chambers with comparable multilayering. In both conditions, the timing of germline-cell death was delayed resulting in intact stage 10 egg chambers. However, these egg chambers continued to exhibit border-cell migration defects, indicating that cnc-KD or Keap1-KD did not rescue the effects of polarity-loss induced by Lgl-KD (Fig.5A, arrowheads).

**Figure 5:**
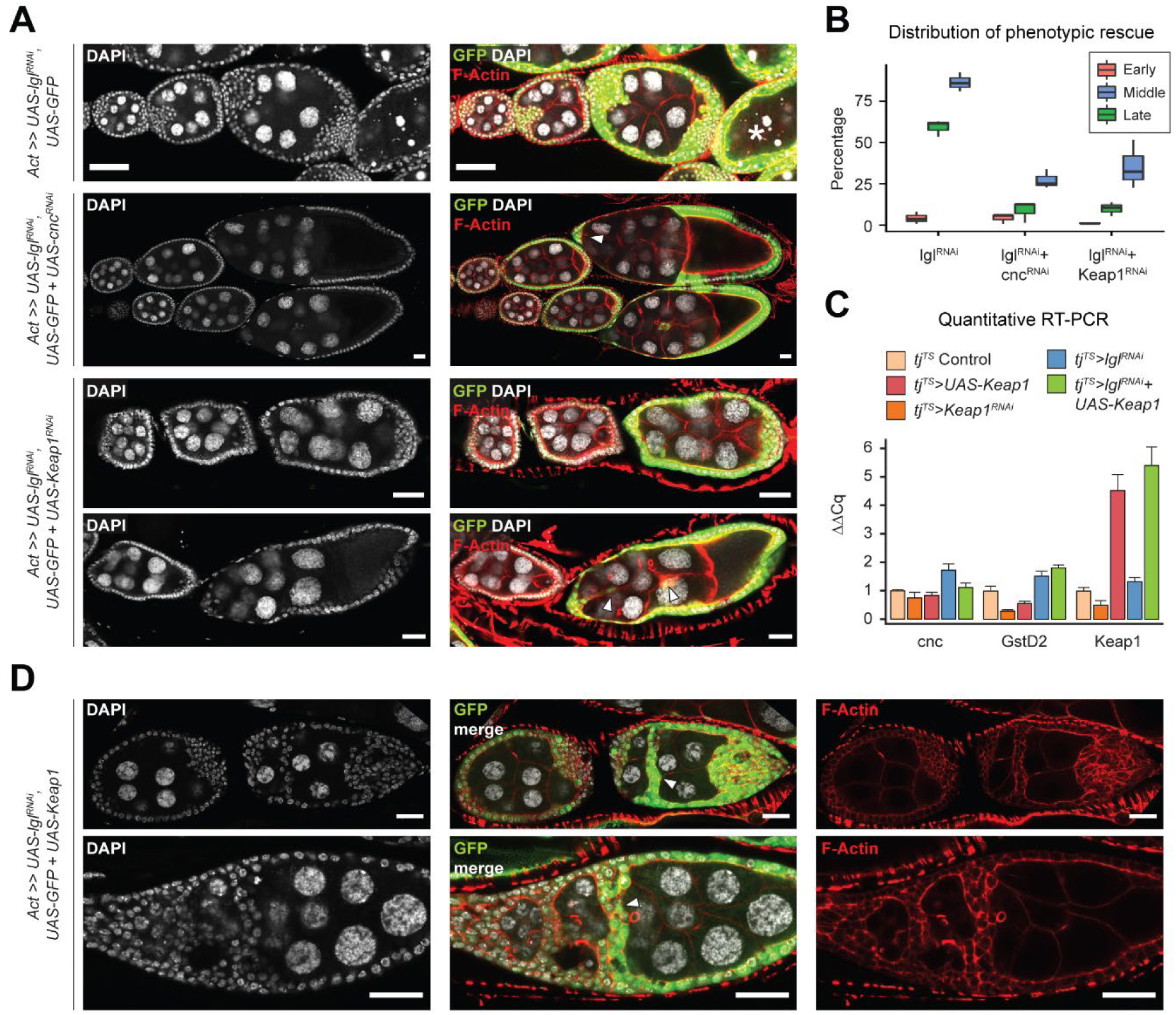
Keap1-Nrf2 signaling is required for Lgl-KD multilayering. **A.** Representative confocal images of ovarioles containing egg chambers with transgene expression in follicle cells (green) driving Lgl-KD (top), Lgl-KD+cnc-KD (middle) and Lgl- KD+Keap1-KD (bottom panels). Degenerating egg chambers are marked by asterisks (*). Border-cell migration defects are indicated by arrowheads. Nucleus is marked by DAPI (white). F-Actin is marked by Phalloidin (red). Scale bars: 20 µm. **B.** Box and Whisker Plot showing the quantification of phenotypic rescue at early (red), mid (blue) and late (green) oogenesis. For the multilayering phenotype at midoogenesis, we have only counted instances of > 2 follicular layers. **C.** Bar Plot showing relative expression of genes involved in the Keap1-Nrf2 signaling pathway. Samples are color-coded as shown in legend. **D.** Confocal images of ovarioles with Lgl-KD+Keap1-OE in follicle cells (green). Arrowheads mark the epithelial bridging phenotype. Nucleus is marked by DAPI (white). F-Actin is marked by Phalloidin (red). Scale bars, 20 µm.

Under normal conditions of redox homeostasis, Keap1 has been shown to target Cnc for ubiquitination and so, Keap1 loss-of-function is expected to promote increased nuclear translocation of Cnc (Itoh et al., 1999). However, contrary to our expectations, genetic epistasis experiments involving *cnc* and *Keap1* knockdowns resulted in comparable phenotypes. To test how *Keap1* manipulation regulates the Keap1-Nrf2 signaling in the ovaries with and without Lgl- KD, we performed quantitative RT-PCR (qRT-PCR) to detect the relative enrichment of pathway components upon *Keap1* loss and overexpression (Fig.5C). We found that in the absence of Lgl-KD, both overexpression and knockdown of *Keap1* did not affect *cnc* expression. On the contrary, manipulating *Keap1* expression in ovaries without Lgl-KD resulted in the decreased expression of the downstream pathway target *GstD2*, which was found enriched specifically in the border cells (Supplementary Fig. 5). Ectopic overexpression (OE) of Keap1 in presence of Lgl-KD (Lgl-KD+Keap1-OE) resulted in a significant increase of Keap1-Nrf2 signaling activity, as confirmed by the levels of *GstD2* being higher than that in Lgl-KD alone, while *cnc* expression was maintained at levels comparable to those observed in the control. In these Lgl-KD+Keap1- OE egg chambers, we noted the worsening of the Lgl-KD phenotype as the egg chambers appeared both multilayered as well as increasingly fused (Fig.5D). We also observed epithelial “bridging” in several egg chambers (n=119/185) where the multilayered cells formed invasive inroads through the recesses between germline cells that connected to the lateral epithelia, forming a bridge between two separate sections of the follicular epithelia (Fig.5D, arrowheads). Collectively, our results suggested that Keap1-Nrf2 signaling directly influenced the behavior of multilayered epithelia, where manipulating *Keap1* expression increased the invasiveness of Lgl- KD cells allowing the formation of bridges between distinct parts of the epithelial layer.

### Elevated *Keap1* increases collective invasiveness of multilayered cells

We next sought to describe the *Keap1* driven enhancement of Lgl-KD cell delamination by assessing cell-cell adhesion as well as actin dynamics. When Lgl-KD+Keap1-OE egg chambers were stained with antibodies against junctional proteins Shg (E-Cad) and Arm (β-Cat), a relative decrease in their enrichment at the multilayered-cell junctions (compared to that in the monolayer) was detected (Fig.6A). However, unlike the apical-most layer of Lgl-KD cells, the delaminated Lgl-KD+Keap1-OE cells did not exhibit a complete loss of cell-cell adhesion, as continued enrichment of Shg and Arm was detected at cell-cell contacts. This observation implied that the cells maintained cellular junctions and likely moved collectively.

**Figure 6:**
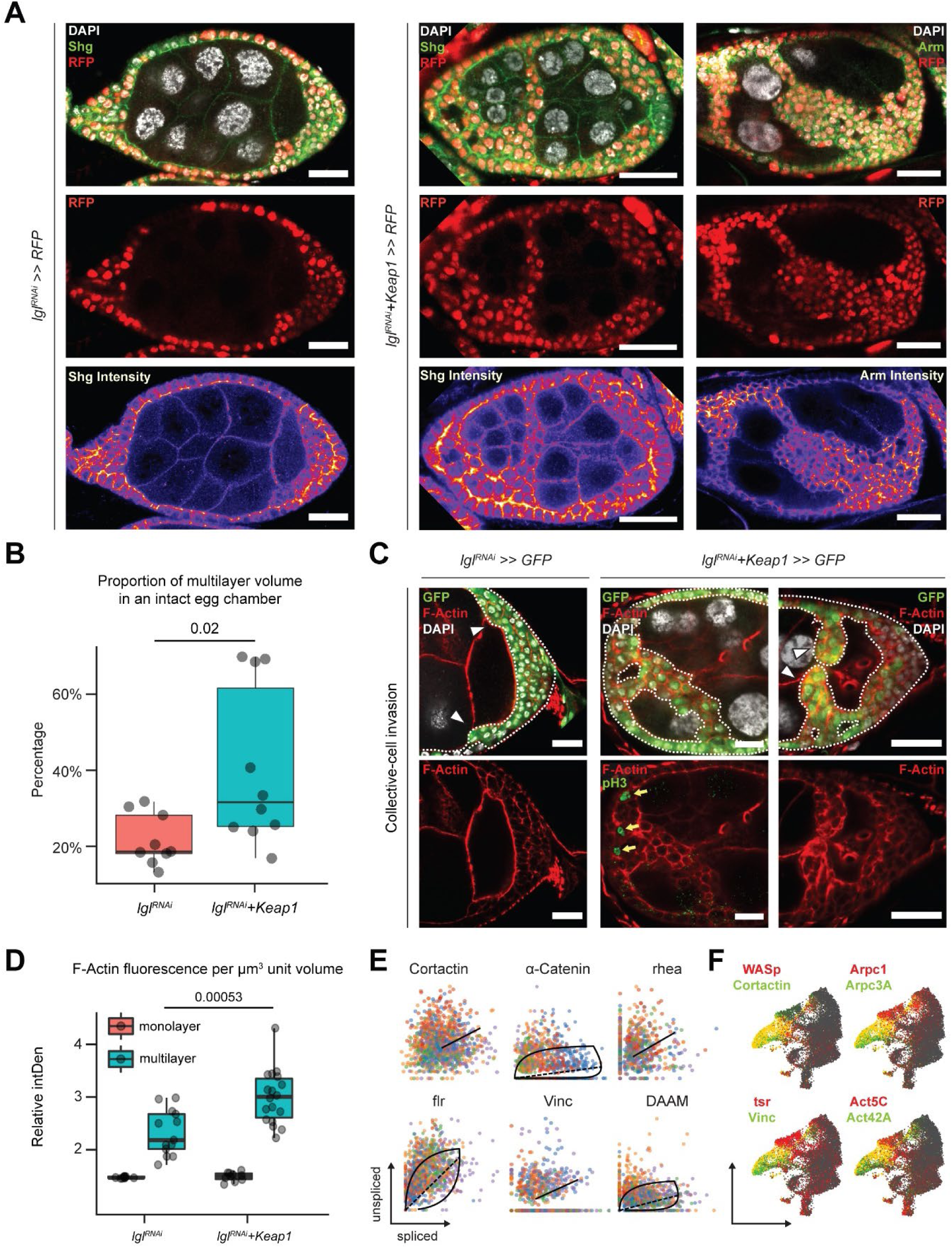
Overexpressing Keap1 in *lgl^RNAi^* follicle cells promote collective invasion of cells via cytoskeletal remodeling. **A.** Confocal images of egg chambers with Lgl-KD (Left) and Lgl-KD+Keap1-OE (middle and right panels) in the follicle cells (red; shown in the middle row). Egg chambers are stained with Shg (DE-Cad; left and center columns) and Arm (right column) (green). Shg and Arm staining intensity is shown in the bottom row. Nucleus is marked by DAPI (white). F-Actin is marked by Phalloidin (red). Scale bars, 20 µm. **B.** Box and Whisker Plot showing the proportion of delaminated epithelial volume compared to the volume of the entire egg chamber. **C.** Confocal images showing collectively invading follicle cells (green) in Lgl-KD (left) and Lgl-KD+Keap1-OE (middle and right). Nucleus is marked by DAPI (white). F-Actin is marked by Phalloidin (red). The invasive fronts are marked by arrowheads. Positive pH3 staining (green in bottom middle panel) shows mitotically dividing cells, marked by yellow arrows. Scale bars, 20 µm. **D.** Box and Whisker Plot to show the relative intensities of F-Actin in the monolayer (red) and multilayers (teal). Individual egg chambers are shown as individual dots. **E.** Phase portraits of cytoskeletal genes in cluster 7. Straight line indicates that the gene exhibits steady state dynamics, while curved solid line with dashed straight line represents upregulation. Individual cells are colored according to their subcluster identity. **F.** Gene enrichment plot showing overlapping enrichment of actin remodeling genes in cluster 7.

Next, we measured the relative proportion of delaminated epithelial volume in total volume of intact egg chambers containing Lgl-KD and Lgl-KD+Keap1-OE in follicle cells. We found that elevating *Keap1* in Lgl-KD background induced a significant (*p=*0.02) but variable increase (40.3%; N=10 egg-chambers at stage 8) in the volume of delaminated epithelial cells compared to Lgl-KD (21.8%; N=9) alone (Fig.6B). The frequent occurrence of holes and gaps within the delaminated epithelia of Lgl-KD+Keap1-OE egg chambers (Fig.5D, 6A and 6C) suggested that the multilayered cells possibly did not expand evenly, as uniform growth would tend to fill all available space. Indeed, mitotic marker phospho-Histone 3 (pH3) was observed only in few cells within the multilayered epithelia of egg chambers with Lgl-KD+Keap1-OE in follicle cells, and was rarely detected inside the invasive front (Fig.6C, middle panel). These observations therefore suggested that the increase in delaminated epithelial volume was not caused primarily by overproliferating cells pushing out the delaminating epithelia.

We subsequently assessed whether the collectively-moving cells display polarized invasion or multicellular streaming by observing F-actin distribution in the invasive cells (Friedl et al., 2012). While F-actin was generally found elevated in the delaminated epithelial bridges formed by Lgl-KD+Keap1-OE cells, when invasive multilayers displaying incomplete bridging were specifically observed, we noticed significantly increased F-actin staining at the leading edge formed by blunt multicellular tips (Fig.6C). Quantifying the relative F-actin intensity in the delaminated epithelia of both Lgl-KD (N=13 invasive multilayers from 10 total egg chambers) and Lgl-KD+Keap1-OE (N=18) egg chambers at midoogenesis revealed: (1) an increase in F-actin enrichment in the multilayered epithelia compared to the monolayer; and (2) a highly-significant (*p=*0.00053) enhancement of F-actin intensity in the Lgl-KD+Keap1-OE multilayers, compared with those in Lgl-KD (Fig.6D). The presence of a defined actin-rich leading edge with multiple cells at the tip indicated that the cells undergo polarized collective invasion.

Collectively invading cells exhibit rapid turnover of cytoskeletal proteins, as well as invadipodial structures in the leading edge. In our single-cell RNA-Seq analysis, we detected dynamic behavior of several genes coding for proteins that mediate interactions between Integrins and filamentous-actin (*e.g., α-Catenin*, *Vinculin* and *cofilin*) (Eddy et al., 2017) and those that regulate actin polymerization (Molnár et al., 2014; Poukkula et al., 2014; Krueger et al., 2019) (such as *Cortactin*, *flr* and *DAAM*, etc.) in cluster 7 cells (Fig.6E and Supplementary Data 6). These genes have known functions in focal adhesion and are required for the formation and maintenance of invadopodia (Aughey, Grice and Liu, 2016; Eddy et al., 2017). We also detected enrichment of genes that support leading-edge protrusion (Eddy et al., 2017) such as *WASp*, *Cortactin*, *Vinculin*, *tsr* (*Drosophila* Cofilin), individual actin molecules (*Act5C* and *Act42A*), and specific genes of the Arp2/3 actin-nucleator complex (*Arpc3A* and *Arpc1*) (Fig.6F). In summary, increasing Keap1 expression promoted collective invasion of the delaminated Lgl- KD cells that exhibit leading edge behavior, possibly by extensive remodeling of the actin cytoskeleton.

## DISCUSSION

Our purpose in this study was to identify genes that regulate the changes in cell plasticity upon apicobasal-polarity loss in epithelial cells, which is arguably one of the earliest characteristics of tumor cells (Moreno-Bueno, Portillo and Cano, 2008; Royer and Lu, 2011). We applied the single-cell RNA-Sequencing (scRNA-Seq) technique followed by marker validation to discover gene expression specific to the distinct phenotypes observed in *Drosophila* ovaries when polarity-gene *lgl* is functionally knocked down in the follicle cells. Identifying cell states that deviate from the normal follicle-cell developmental lineage upon Lgl loss-of-function, we revealed their transcriptomic signatures and underlying regulon activity at an unprecedented resolution. Our analyses highlight a complex transcriptional ecosystem where a network of regulons function with varying degrees of specificity in the heterogenous cells of the delaminating epithelia. This study represents an important step toward systematically describing the polarity-loss induced tumor model in the *Drosophila* follicle cells to understand intra-tumor expression heterogeneity and characterize the changes to cell plasticity following polarity loss.

Among our primary findings is the identification of a heterogenous group of cells that are characterized by an enrichment of tumorigenic-stress signaling (Bunker et al., 2015; Mundorf et al., 2019; Hamaratoglu and Atkins, 2020). Amenable to targeted analysis, scRNA-Seq allowed us to further classify this cluster, enabling the collective analysis of cells at contrasting fates as well as states of stable transcription. Modeling the inherent RNA velocity in each cell, we inferred the lineage relationships among the heterogenous population of cells. Supported by the observation that mature post-mitotic cells form the invasive front of delaminating multilayers, we specifically focused on cells that transitioned out of the post-mitotic follicle-cell cluster and exhibited an upregulation of Keap1-Nrf2 signaling. Keap1-Nrf2 signaling is activated in these cells likely as a response to elevated ER stress (evidenced by an enrichment of *crc*, *Xbp1* and several CREB/ATF transcription-factors associated regulons) as has been previously reported (Wortel et al., 2017), and not on account of significant oxidative stress as neither the superoxide-detector dihydroethidium (DHE) nor the common oxidative-stress response genes were detected in the multilayers or in cluster 7 respectively (Supplementary Fig. 6). However, the role of oxidative stress cannot be ruled out entirely since a stressed ER can also cause reactive oxygen species (ROS) accumulation (Wortel et al., 2017; Limia et al., 2019) that would directly impact Keap1 structure thereby blocking Nrf2-mediated transcription, which is not the case in Lgl-KD samples. Considering this redundancy in signaling regulation, it is difficult to ascertain the direct cause of Keap1-Nrf2 pathway activation in our model from these experiments alone. Our conclusions from this study therefore warrant further investigations to understand the precise cause of Keap1-Nrf2 signaling activation in the cells with polarity loss.

Under conditions of homeostasis, Cnc (*Drosophila* homolog of Nrf2) binds to its endogenous inhibitor Keap1 in the cytoplasm that negatively regulates Cnc-driven transcription of many antioxidant (Sykiotis and Bohmann, 2008) and anti-inflammatory enzymes (Warren L. Wu and Papagiannakopoulos, 2020). In our study, inducing loss-of-function of both *Keap1* and *cnc* in cells with Lgl knockdown (Lgl-KD) reduced the severe multilayering phenotype. This identical outcome of phenotypic rescue was unexpected given the antagonizing functional relationship between the proteins. Since apoptosis was rarely observed in these cells (Dcp-1 data not shown), the apparent loss of multilayered cells was likely not a consequence of gene-knockdown associated cytotoxicity, therefore demanding an alternate explanation for the apparent rescue. Ectopic expression of Keap1, on the other hand, induced increased invasiveness of the delaminated cells, thereby suggesting that its loss likely rescued the invasive behavior of the multilayers in particular. Outside its more commonly-defined role in oxidative-stress response, cell-culture based studies have indeed identified additional roles of Keap1 in regulating invasive-cell behavior (Rachakonda et al., 2010) and maintaining cytoskeletal stability (Kang et al., 2004; Yamaguchi and Condeelis, 2007). It is also hypothesized that *wild-type* Keap1 may stabilize focal adhesion-like assemblies to regulate the cytoskeleton (Velichkova and Hasson, 2003; Dinkova-Kostova, Holtzclaw and Kensler, 2005), which might partially play a role in the collectively-invading Lgl-KD cells observed in our study. Given that the role of Keap1-Nrf2 signaling in cancers is more commonly investigated by using loss-of-function mutants of *KEAP1* (Warren L Wu and Papagiannakopoulos, 2020), the role of its *wild-type* protein in cancer remain largely unexplored. Our study, therefore, ascribes a highly relevant function to *wild-type* Keap1 that it stands to play at the earliest stages of tumorigenesis when the cellular burden of mutations is low (Bozic et al., 2010) and dysplastic cells exhibit loss of polarity (Thiery, 2003). Recently, dynamic regulation of Nrf2 within the collectively-invading cancer cells has been shown to also have an effect on the metastable states of cancer cells displaying partial-EMT (Zhou et al., 2016; Bocci et al., 2019; Vilchez Mercedes et al., 2022). It is therefore also possible that in our experiments, Keap1 expression regulates the metastability of Lgl-KD cells indirectly, as a result of which, the invasive behavior of the delaminated tissue on the whole is being determined.

Enriched F-actin at the leading edge of invasive fronts is a hallmark of invasive membrane protrusions of leader cells in a collectively migrating cell cohort (Eddy et al., 2017) and is frequently observed in tumor buds (Grigore et al., 2016). We were able to characterize the behavior of collectively-migrating Lgl-KD cells also expressing Keap1 by staining for junctional and cytoskeletal markers (Friedl et al., 2012), and concluded that the delaminated cells showed leading edge behavior based on the differential enrichment of F-actin in the leading cells. Moreover, the presence of holes within the epithelia – which is not observed upon polarity-gene loss-of-function alone (Goode, Wei and Kishore, 2005; Szafranski and Goode, 2007) – further suggested that the delaminating cells exhibit a differential capacity to migrate, as the holes would likely have been filled if every cell could either exert independent migratory force or exhibit uniform proliferation to compensate for the increased invasiness. It is thus likely that the increase in polarized invasiveness is able to outrun the pace of cell division, leaving behind gaps in the delaminated epithelia. Despite strong evidence suggesting that ectopic Keap1 drives leading-edge directed collective invasion of Lgl-KD multilayers, our conclusions are mildly tempered by the constraints of available space within the egg chambers which limits the ability to separate collective invasion maintained by weak cell-cell adhesions and random movements of cells in the narrow passage between the germline cells. Time-lapse or live imaging of this invasive behavior could be used in subsequent studies to understand the underlying mechanism better. Nonetheless, our approach in this study presents an innovative analytical framework enabling the high-resolution characterization of cells that separate from the expected lineage, applying which, novel regulators of tumor-cell invasiveness were identified in the *Drosophila* follicle cells upon polarity loss.

## Methods

### Fly husbandry

Flies were reared under standard lab conditions at 25°C and were fed dry yeast a day before dissections. For temperature-sensitive RNAi experiments, *tj-Gal4* driver was combined with *tub- Gal80^ts^* and the resulting flies were reared at 18°C to repress spurious Gal4 activity and transferred to 29°C for transgene induction. These flies were kept at 29°C for the required number of days, following which, they were dissected. Combining *tj-Gal4, UAS-GFP, tub- Gal80^ts^* with *UAS-l(2)gl^RNAi^, UAS-Dcr2* resulted in increased genetic-lethality and only few flies were recovered. Therefore, *tj-Gal4* lines without a fluorescence-reporter were used to generate a stable line and the resultant phenotype was tested for comparable results. For FLPout experiments, flies were reared at 25°C and were heat-shocked at 37°C for 20 minutes, fed with dry yeast 2 days after heat-shock (AHS) and were dissected 3 days AHS.

### Fly stocks

BDSC stocks:

*UAS-Lgl-RNAi*: y[1] v[1]; P{y[+t7.7] v[+t1.8]=TRiP.HMS01905}attP40 (#38989),

*UAS-Keap1*: y[1] w[67c23]; P{y[+mDint2] w[+mC]=EPgy2}Keap1[EY02632] (#15427),

*UAS-Keap1-RNAi*: y[1] sc[*] v[1] sev[21]; P{y[+t7.7] v[+t1.8]=TRiP.HMS02180}attP40 (#40932),

*UAS-cnc-RNAi*: y[1] v[1]; P{y[+t7.7] v[+t1.8]=TRiP.JF02006}attP2 (#25984),

*hsFLP*: P{ry[+t7.2]=hsFLP}1, y[1] w[1118]; Dr[Mio]/TM3, ry[*] Sb[1] (#7),

*Act>y>Gal4, UAS-GFP*: y[1] w[*]; P{w[+mC]=AyGAL4}25 P{w[+mC]=UAS-GFP.S65T}Myo31DF[T2] (#4411).

*hsFLP;; Act>CD2>Gal4, UAS-hRFP*: w[*]; P{ry[+t7.2]=Act5C(FRT.polyA)lacZ.nls1}2,

P{w[+mC]=Ubi-p63E(FRT.STOP)Stinger}9F6/CyO; P{w[+mC]=GAL4-Act5C(FRT.CD2).P}S,

P{w[+mC]=UAS-His-RFP}3/TM3, Sb[1] (modified from #51308).

VDRC stocks:

*UAS-l(2)gl-RNAi* (#51247).

Kyoto stocks:

*tj-Gal4*: y[*] w[*]; P{w[+mW.hs]=GawB}NP1624/CyO, P{w[-]=UAS-lacZ.UW14}UW14 (#104055).

*Slbo-GAL4* (gift from Dr. D. Montell).

*GstD:lacZ* (gift from Dr. D. Bohmann).

*UAS-Pxn* (gift from Dr. U. Banerjee).

### Immunofluorescence Staining, Imaging, and Figure Preparation

Flies were dissected at room temperature in 1X PBS (Phosphate-Buffered Saline), and the ovaries were fixed for 15 minutes in 4% PFA (paraformaldehyde). Fixed ovaries were then washed 3 times in 1X PBT (PBS with 0.2% Triton X-100) for 20 minutes per wash and were incubated for 1 hour with blocking solution (1X PBT with 0.5% BSA and normal goat serum). Incubation with primary antibody, diluted in blocking solution, was performed overnight at 4°C. The primary antibody-stained ovaries were again washed 3 times with 1X PBT the following day and were subsequently incubated with secondary antibodies diluted in blocking solution for 2 hours at room temperature. After washing again for 3 times with 1X PBT, and once with PBS, the ovaries were dyed with DAPI (Invitrogen, 1µg/mL) to stain the nuclei. Samples were finally mounted on microscopic slides after adding 80% glycerol mounting solution.

The following antibodies or dyes are mentioned in this paper: DSHB:

mouse anti-Arm (N27A1, 1:40 dilution),

mouse anti-Cut (2B10, 1:30),

rat anti-Shg (DCAD2, 1:20),

mouse anti-Hnt (1G9, 1:15),

rat anti-Mmp1 (1:1:1 mixture of 3B8, 3A6 and 5H7, 1:40),

mouse anti-Sn (sn7C, 1:25).

mouse anti-Drpr (5D14, 1:50) Promega:

mouse anti-β-Gal (PAZ3783, 1:500). Millipore:

rabbit anti-pH3 (06-570, 1:200). Invitrogen:

Phalloidin Flour 546 and 633 (A22283 and A22284; 1:50)

Secondary antibodies:

Alexa Fluor 488, 546, and 633 (1:400, Molecular Probes).

For real-time Dihydroethidium (DHE) staining, *tj^TS^>lgl^RNAi^* ovaries (with no UAS-tagged fluorescence reporter) were dissected in Grace’s Insect Basal medium (VWR; #45000-476), and incubated in Dihydroethidium (Invitrogen, D1168; diluted in 0.1 µmol/L DMSO) for 5 minutes.

Following DHE incubation, ovaries were fixed and standard protocol for immunofluorescence were followed.

All images were acquired using the Zeiss LSM 800 confocal microscope and the proprietary Zeiss microscope software (ZEN blue). Images comparing different samples under comparable experimental conditions were obtained using the fixed settings for image acquisition. Final images were processed and analyzed using FIJI-ImageJ (Schindelin et al., 2012) open-source software.

Individual images were organized into panels and were annotated in Adobe Illustrator.

### <u>Q</u>uantitative <u>R</u>eal <u>T</u>ime (qRT)-PCR

Entire ovaries were lysed in TRIzol (Invitrogen) and the RNAs were prepared according to the manufacturer’s protocol and quantified using nanodrop. For each sample, 1μg of mRNA was reverse-transcribed using oligo-dT-VN primers and ImProm-II as the reverse transcriptase (Promega) in triplicate. Real-time quantitative amplification of RNA was performed using the Sybr Green qPCR Super Mix (Invitrogen) and the iQ™5 Real-Time PCR Detection System from BioRad according to the manufacturer’s protocol. The relative expression of indicated transcripts was quantified with the CFX_Manager Software (BioRad) using the 2[−ΔΔC(T)] method. According to this method, the C(T) values for the expression of each transcript in each sample were normalized to the C(T) values of the control mRNA (*GAPDH*) in the same sample. The values of untreated cell samples were then set to 100% and the percentage transcript expression was calculated.

The following primer pairs have been used in this paper:

Ets21C-F: GAGGCCGATTAATGCCATGC

Ets21C-R: AGTTGAGGGCGGGTAATTGG

grnd-F: TCGGTCAGGAAGTTGAGTGC

grnd-R: CGCACAGAAACGCATCGTAG

Jra-F: AACACATCCACCCCGAATCC

Jra-R: CCTTGGTGGGGAACACCTTT

kay-F: TTTCTGCCCGCCGATCTAAG

kay-R: GTTGCCGAGGATAAGATTGCG

Mmp1-F: CAAGTTGGACGAGGACGACA

Mmp1-R: GTAGGCCTCAGCTGGTTTGT

cnc-F: CCACAACACCACCGGGAATA

cnc-R: ATGTGGCGTGAGGAAAGTGT

GstD2-F: CCGGATCGGATGAGGACTTG

GstD2-R: TTCGAACGTGGAGACAGTGG

Keap1-F: TTCCTGCAGCTTTCGGCATA

Keap1-R: GCTCCTCCTGCACATTCAGT

### Bulk RNA-Sequencing Sample Preparation for Sequencing

Flies of the genotypes *tj>GAL4^TS^, UAS-GFP* (control) and *tj>GAL4^TS^, UAS-l(2)gl^RNAi^* (no GFP) (experiment) were raised in 18°C for 2-3 days post-eclosion. They were then transferred to 29°C for either 24 hours (1 day) or 96 hours (4 days) to induce the multilayering of follicle cells for short and long periods of time. All female flies had access to males in their vials and were given yeast supplement 24 hours prior to dissection to facilitate oogenesis. Ovaries were dissected from 40 flies in Complete medium (Grace’s Insect Basal medium (VWR; #45000-476) supplemented with 15% FBS (ATCC; #30-2020). The bulk of the ovary was severed from the oviduct, fully developed eggs were crudely removed and individual ovarioles were roughly separated from each other. Each sample was then transferred to a sterile microfuge tube and the media was aspirated. The remaining ovaries of each sample was then flash-frozen in liquid Nitrogen and were stored in -80°C prior to library preparation.

Total RNA libraries were made using the NEBNext Ultra II Directional RNA library Prep Kit for Illumina using the established protocol for use with NEBNext Poly(A) mRNA Magnetic Isolation Module (NEB#E7490). We used Rapid Run OBCG single read 50 bp on the Illumina Hi-Seq 2500 system to sequence these libraries with 2 technical replicates for each time-point and genotype. Sequenced reads were demultiplexed and the indexes were removed using CASAVA v1.8.2 (Illumina).

### Single-Cell RNA Sequencing Sample Preparation

*tj^TS^>lgl^RNAi^* flies were shifted to 29°C, permissive temperature for the activation of transgenic expression of short-hairpins targeting the *l(2)gl* mRNA for functional knockdown in every cell of the follicular lineage. Ovaries from 50 such adult female flies were dissected 72 hours (3 days) later in Complete medium and were then transferred to a tube containing 300 µl EBSS (TF, #78154; no calcium, magnesium, and phenol red) and were gently mixed for 2 minutes. After allowing the tissue to settle, EBSS was removed and individual cells were dissociated from the tissue in 100 µl Papain (Worthington, #LK03178; 50 U/mL in EBSS and heat activated in 37°C for 15 minutes before use) for 30 minutes. The suspension was mechanically dissociated every 3 minutes by gentle pipetting up and down. To quench the digestion, 500 µl Complete medium was subsequently added to the dissociated cells. The suspension was then passed through a 40 µl sterile cell strainer and centrifuged for 10 minutes at 700 RCF. The supernatant was then removed and 100 µl of Complete medium was added. Viability and concentration of cells were assayed using Propidium Iodide staining and manual cell count estimates were made using a hemocytometer to adjust input cell concentration for single-cell RNA-Seq library preparation.

### 10X Genomics Library Preparation and Sequencing

Single-cell libraries were prepared using the Single Cell 3’ Library & Gel Bead Kit v2 and Chip Kit (10X Genomics; PN-120237) according to the recommended 10X Genomics protocol and the single cell suspension was subsequently loaded on to the Chromium Controller (10X Genomics). Library quantification assays and quality check analysis were performed using the 2100 Bioanalyzer instrument (Agilent Technologies). The library samples were then diluted to the final concentration of 10nM and loaded onto two lanes of the NovaSeq 6000 (Illumina) instrument flow-cell for a 100-cycle sequencing run. Reads were demultiplexed the same way as described in Bulk RNA-Sequencing sample preparation and sequencing and fastq files were generated for each library that were subsequently processed in the Cellranger CLI software (10X Genomics).

### Statistics And Reproducibility

All statistical analysis of data and generation of graphs were performed in R. Boxes in box plots show the median and interquartile range; lines show the range of values within 1.5× of the interquartile range. Student’s t-test has been used to perform statistical analyses. All phenotypic observations are independently assessed by two individuals, where one was blinded to the experimental goal (but aware of the experimental conditions). All images are representative of at least three independent experiments. Each experiment involves the dissection of a minimum of 25 flies per genetic cross. The genotypes for each experiment are clearly mentioned in each figure.

The scRNA-Seq analysis of the *w^1118^* ovarian cells in this study reused the sequences generated for the published *w^1118^* ovarian cell-atlas (Jevitt et al., 2020), that were obtained from the SRA database (SRX7814226). The number of cells passing the filtering criteria of Cellranger increased significantly from the previously-reported number of 12,671 cells with 28,995 mean reads-per-cell to 24,144 cells with 17,095 mean reads-per-cell, with no change in the total number of sequenced reads (429,855,892). This result was a result of re-mapping of sequences to the top-level dm6 reference genome, which was different from the reference genome used in the Jevitt et al. study. This sequence realignment improved the genome alignment of the reads. The adopted analysis pipeline in this study, however, does not change the primary findings of the original paper as the markers described for each cell-types remain unchanged.

Gene enrichment plots for scRNA-Seq datasets were scaled within the range of 5^th^ and 95^th^ quartiles to increase cluster-specific enrichment of gene expression.

### Bulk RNA Sequencing Analysis

Quality control on the sequences is performed using FastQC v0.11.9. Reads are mapped to the *Drosophila melanogaster* Release-6 (BDGP6.28.dna.toplevel) reference genome using STAR v2.7.0a (Dobin et al., 2013). BAM files resulting from STAR are sorted using the featureCounts program (http://bioinf.wehi.edu.au/featureCounts/) of the Subread package v2.0.1. Low-abundance counts of < 1 CPM (counts per million) are removed and the downstream processing is performed using the edgeR pipeline 3.28.1 (Robinson, McCarthy and Smyth, 2009) in R for the Principle Component Analysis of the two-sample experiment with two replicates each.

### Single-cell RNA Sequencing Data Processing

#### Cellranger and Velocyto

Raw fastq sequencing files from each of the 10X Genomics Chromium single-cell 3’ RNA-seq libraries were processed using Cellranger (version 3.0.0). The reference index for Cellranger was first built using the *Drosophila melanogaster* Release-6 (BDGP6.28.dna.toplevel) reference genome (ftp://ftp.ensembl.org/pub/release-102/gtf/drosophila_melanogaster/). The Cellranger count pipeline for alignment, filtering, barcode and UMI counting was used to generate the multidimensional feature-barcode matrix for all samples. The Cellranger derived bam file for *tj^TS^>lgl^RNAi^* sample was further processed using Velocyto CLI (default parameters), for the annotations of unspliced/spliced reads (La Manno et al., 2018).

#### Seurat

The R package Seurat v3.0 (Stuart et al., 2019) is a universally popular software program to perform single-cell RNA-Seq data preprocessing, dimensionality reduction, cell clustering analyses, differential gene expression analysis and for general dataset handling. Since the detailed steps of data analyses in Seurat are explained on their website (https://satijalab.org/seurat/), we only describe the schematic workflow applied in this study.

Each sample was filtered for low-quality cells by setting sample-specific thresholds for UMIs, gene counts and mitochondrial gene-expression. We used 1,800 genes per cell as the upper threshold and 650 genes as the lower threshold for the control sample and 1,800 genes per cell and 600 genes as the upper and lower thresholds (respectively) for the *tj^TS^>lgl^RNAi^* sample. Then only the cells having under 12,000 (control) and 15,000 (*tj^TS^>lgl^RNAi^*) UMI counts were selected. Finally, an upper limit for the mitochondrial gene-expression was applied to limit the selection of dying cells by filtering out cells expressing more than 10% (control) and 4.5% (*tj^TS^>lgl^RNAi^*) of genes whose symbols begin with “mt:” that is indicative of mitochondrial genes. As a result, 19,986 cells are finally obtained for control and 16,060 cells for *tj^TS^>lgl^RNAi^* sample and these cells were subjected to downstream analyses. After the cells were embedded on lower UMAP dimensions following the suggested parameters for dimensionality reduction, the follicle cell-lineage (epithelial cell-type in the ovary) was further isolated from both the control and *tj^TS^>lgl^RNAi^* sample by removing irrelevant cell-types, such as the germline cells (marked by the expression of specific markers *vasa* and *osk*), the muscle sheath cells (*Zasp66* and *Mp20*), the oviduct tissue (*abd-A*) and the hemocytes (*Hml*). We did not detect the oviduct tissue in the *tj^TS^>lgl^RNAi^* dataset.

### Integration of scRNA-Seq Datasets and Information Extraction

For the integration of *w^1118^* and *tj^TS^>lgl^RNAi^* follicle cell lineages, cells were aligned using the functions *FindIntegrationAnchors()* and *IntegrateData()* with 2000 genes for anchor finding and 50 dimensions for the Canonical Correlation Analysis (CCA). UMI counts of the “integrated” assay were normalized, log-transformed and scaled, following which the cells were clustered using 60 Principle Components (PCs) and a resolution parameter of *1* across the two datasets. Cells were finally embedded on lower UMAP dimensions, using 100 neighboring points and a minimum distance of 0.5 for local approximations of manifold structure. The integrated dataset was carefully processed with an enforced awareness of the selected assay for downstream analyses. We used Seurat-integrated ALRA imputation (Linderman, Zhao and Kluger, 2018) on the sparse count-matrix in the “spliced” assay to recover missing values for gene count (default settings). The imputed matrix is stored in the “alra” assay and was used only to generate gene-enrichment plots. To identify differential markers, the analysis was restricted to the “spliced” assay.

### RNA velocity analysis

#### scVelo

The loom file generated by Velocyto was used as an input in Seurat-based analysis. Subsetted *tj^TS^>lgl^RNAi^* cells were further passed for lineage inference by applying scVelo (Bergen et al., 2020). Without additionally processing the dataset, we directly estimated the underlying RNA velocity on the Seurat-determined cluster identities. We ran the dynamic model to learn the full transcriptional dynamics of the splicing kinetics in these cells. Velocity pseudotime, underlying latent time and terminal end-points were determined for the selected cells using default parameters. Cluster-specific genes that drive pronounced dynamic behavior were detected and their phase portraits were generated where individual cells are colored according to their subcluster identities in cluster 7.

### Regulon Analysis using SCENIC

Regulon activity was assessed using the R package SCENIC v1.1.2-2 (Aibar, 2017). Expression matrix from the “spliced” assay in each cell along with the corresponding Seurat metadata were used as input for SCENIC. The gene sets forming individual regulons were identified using the cisTarget v8 motif collection dm6-5kb-upstream-full-tx-11species.mc8nr (https://resources.aertslab.org/cistarget/databases/drosophila_melanogaster/dm6/flybase_r6.02/mc8nr/gene_based/dm6-5kb-upstream-full-tx-11species.mc8nr.feather). Overlapping regulon modules were identified based on the Connection Specificity Index (CSI), a context-dependent measure for identifying specific associating partners. CSI matrix was generated using the open-source R package scFunctions (https://rdrr.io/github/FloWuenne/scFunctions/).

### Data availability

Both raw and processed sequencing data is available at GSE175435.

## Acknowledgements

We would like to thank Gengqiang Xie, Sumei Zhang, Roger Mercer, Yanming Yang, Cynthia Vied, Amber Brown, and Brian Washburn (at Florida State University Biological Science Core) for their assistance in library preparation and sequencing. 10X Chromium controller and other essential hardware was provided by the FSU College of Medicine Translational Science Laboratory. We deeply appreciate feedbacks from Shangyu Gong, Chih-Hsuan Chang, Hongcun Bao, Fei Cong, Jie Sun, Calder Ellsworth, Anindita Baruah and Chun-Ming Lai.

Special thanks to Norbert Perrimon Lab, Hugo Bellen Lab, Jin Jiang Lab, and the Gene Disruption Project for their contributions in generating the transgenic lines used in our study.

## Author Contributions

D.C. designed and performed all experiments, analyzed both the wet-lab and computational data and wrote the manuscript. X.W. assisted with the qRT-PCR experiments. C.C.A.M. and Y.H. assisted with fly husbandry. D.C. and A.J. prepared and maintained samples for RNA-Seq experiments. X.W. prepared the RNA-Seq libraries. D.C. and W.D. were involved in manuscript conceptualization. W.D. supervised, acquired funding and administered the study.

## Ethics Declarations

The authors declare no competing interests.

## SUPPLEMENTARY FIGURE LEGENDS

**Supplementary Figure 1:**
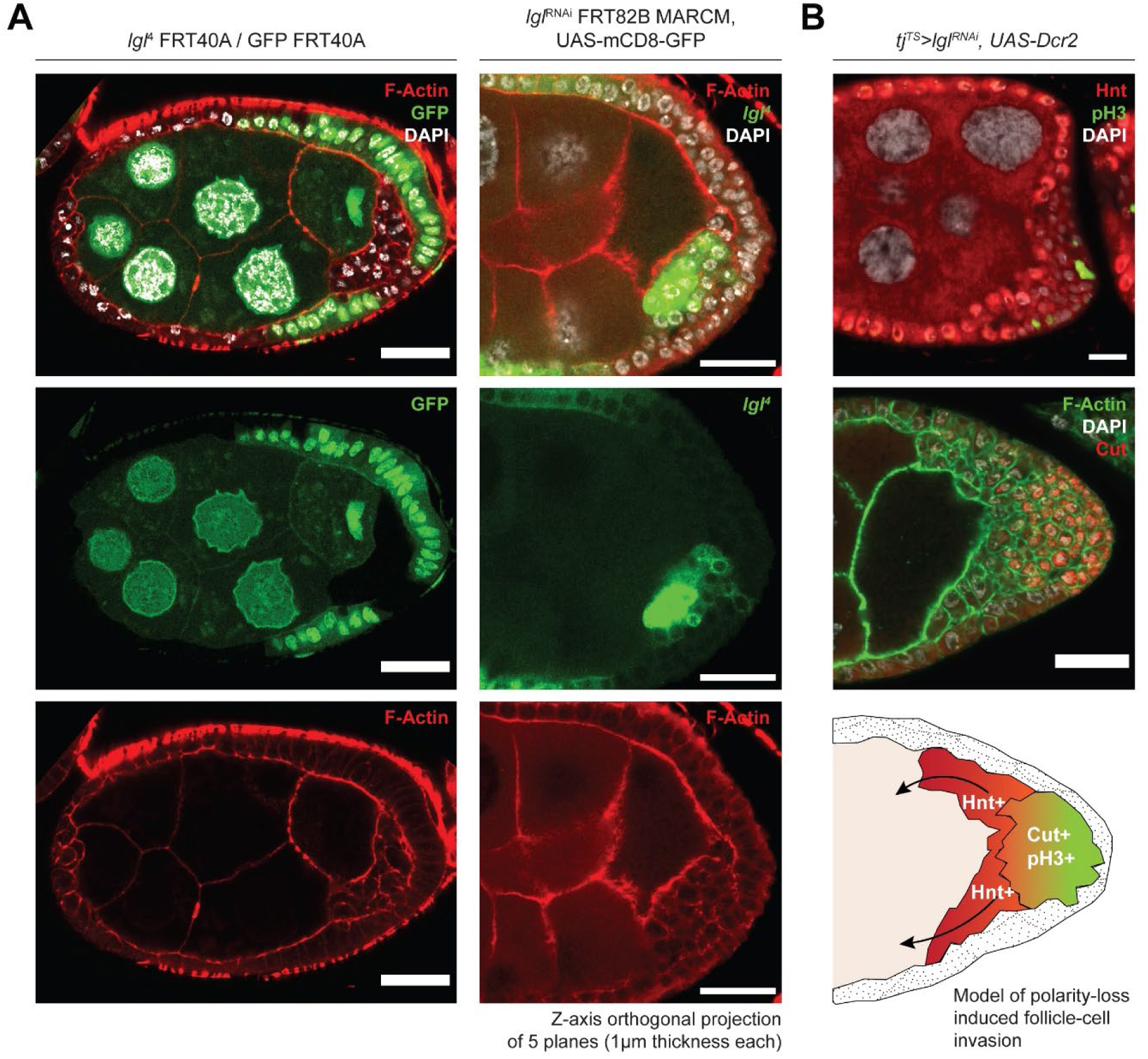
Lgl loss-of-function in follicle cells causes invasive multilayering showing cell-fate heterogeneity. **A.** Confocal Images of egg chambers containing mitotic clones (GFP-) of *lgl^4^/lgl^4^* follicle cells (Left) and MARCM clones (GFP+) of *lgl^RNAi^/lgl^RNAi^* follicle cells (Right). GFP is colored in green, Nucleus is marked by DAPI (white) and F-Actin is marked by Phalloidin (red). **B.** Confocal Images showing common cell-fate markers Hnt, pH3 (top panel) and Cut (lower panel). Nucleus is marked by DAPI (white). Scale bars, 20 µm. A model of apically-invasive multilayers exhibiting cell-fate heterogeneity is shown in the bottom-right panel.

**Supplementary Figure 2:**
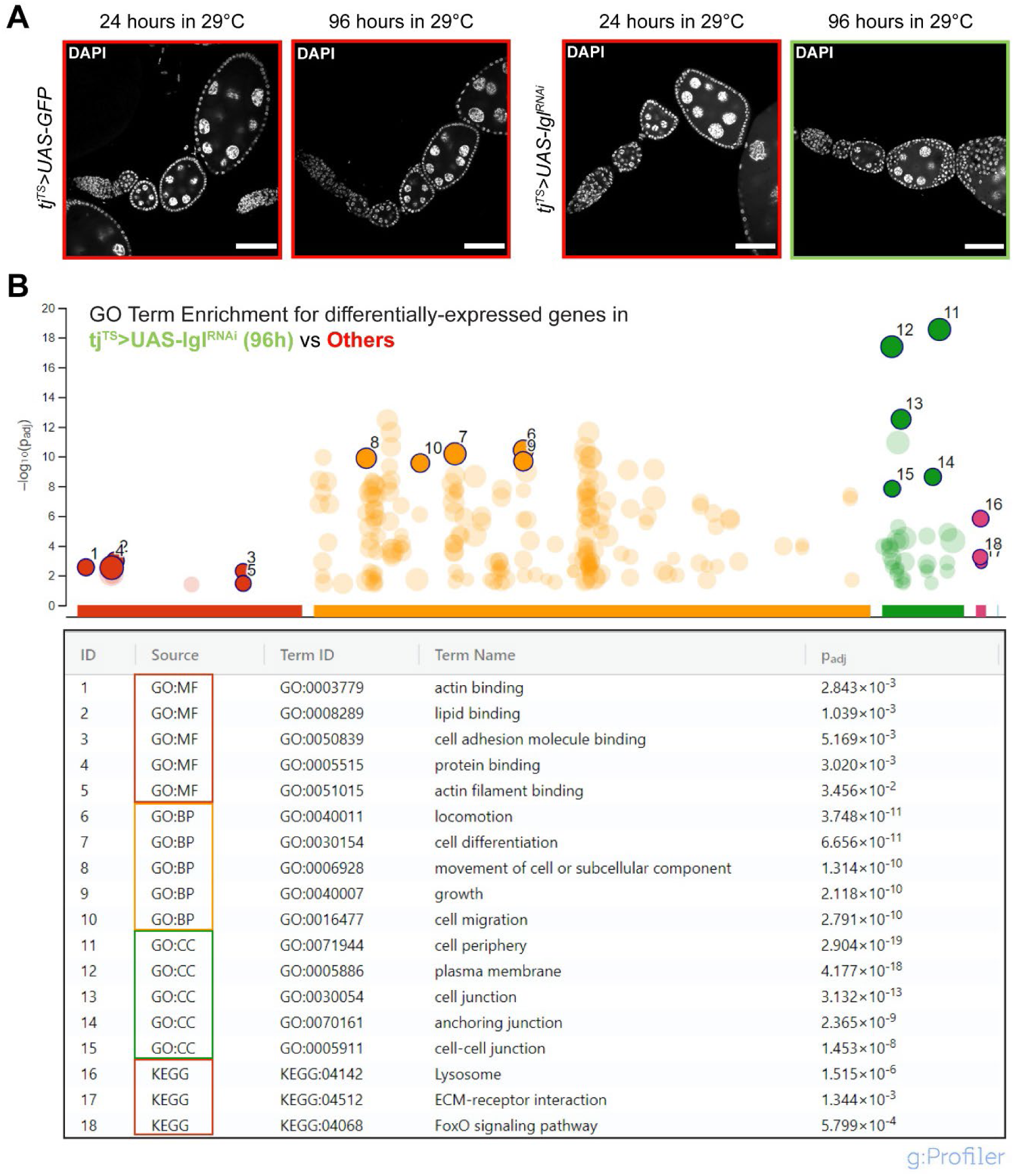
Whole-tissue RNA-Seq of samples containing multilayered Lgl- KD follicle cells. **A.** Representative confocal images of samples used as input for RNA-Seq analysis. **B.** Enriched GO Terms in the 477 differentially-expressed genes found by comparing 96h-Lgl-KD samples (marked by green border in A) with the rest of the samples (marked by red borders in A). The highlighted GO Terms are mentioned in the table below with their corresponding adjusted p-values. Scale bars, 50 µm. See also Supplementary Data 1.

**Supplementary Figure 3:**
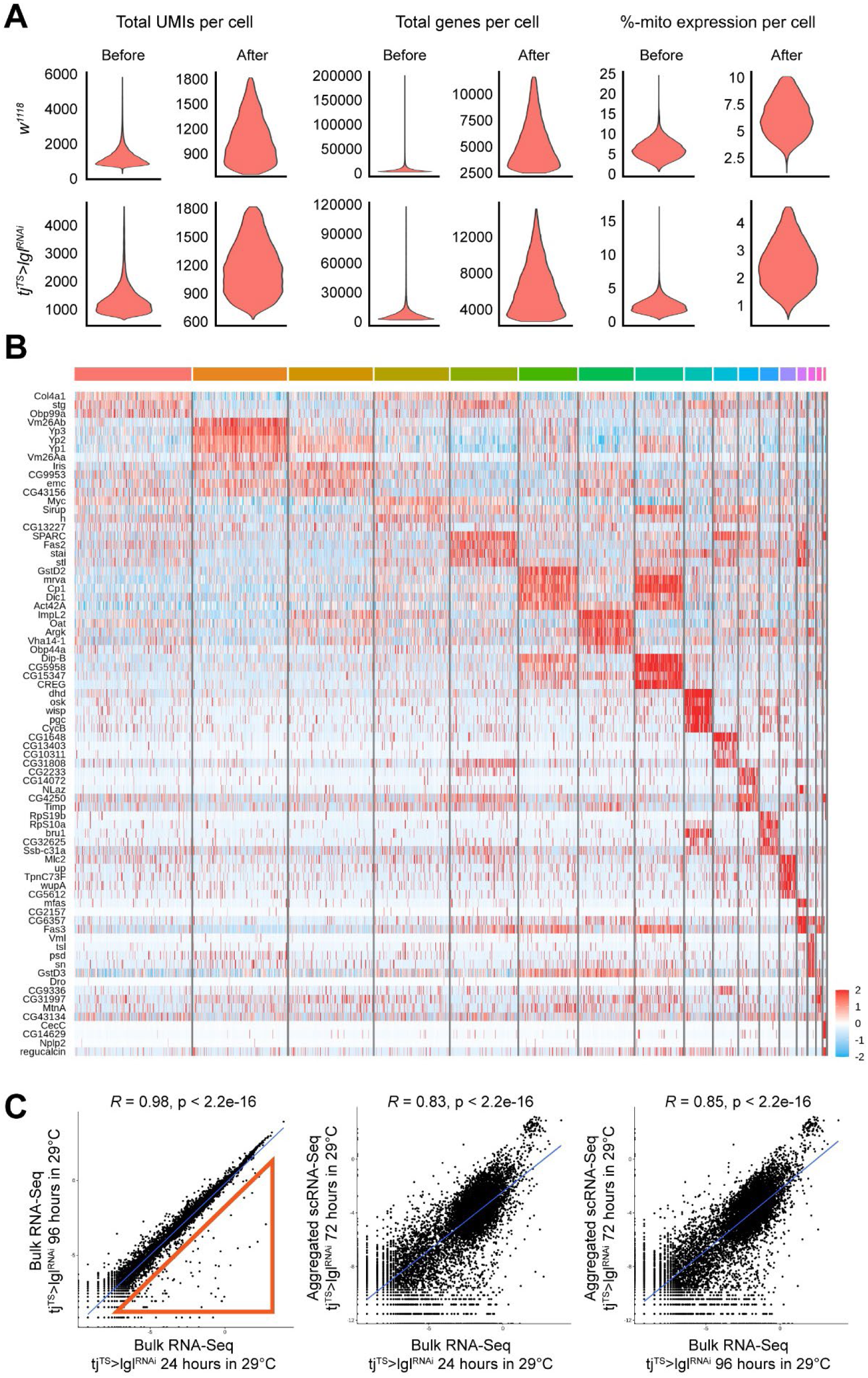
Single-Cell RNA-Seq of ovaries with 72h-Lgl-KD follicle cells. **A.** Violin Plots showing the distribution of total UMI per cell (left column), total genes per cell (middle column) and percentage of cells expressing mitochondrial genes per cell (right column) before and after filtration. Top row represents isolated *w^1118^* cells while bottom row represents *tj^TS^>lgl^RNAi^* cells. **B.** Heatmap of cluster-specific markers for individual *tj^TS^>lgl^RNAi^* clusters shown in Fig.1F. **C.** Correlation scatter plots to show the similarities between RNA-Seq samples. Red triangle in the left plot represents late-stage follicle-cell lineage markers indicating late-stage developmental failure in 96h-Lgl-KD samples.

**Supplementary Figure 4:**
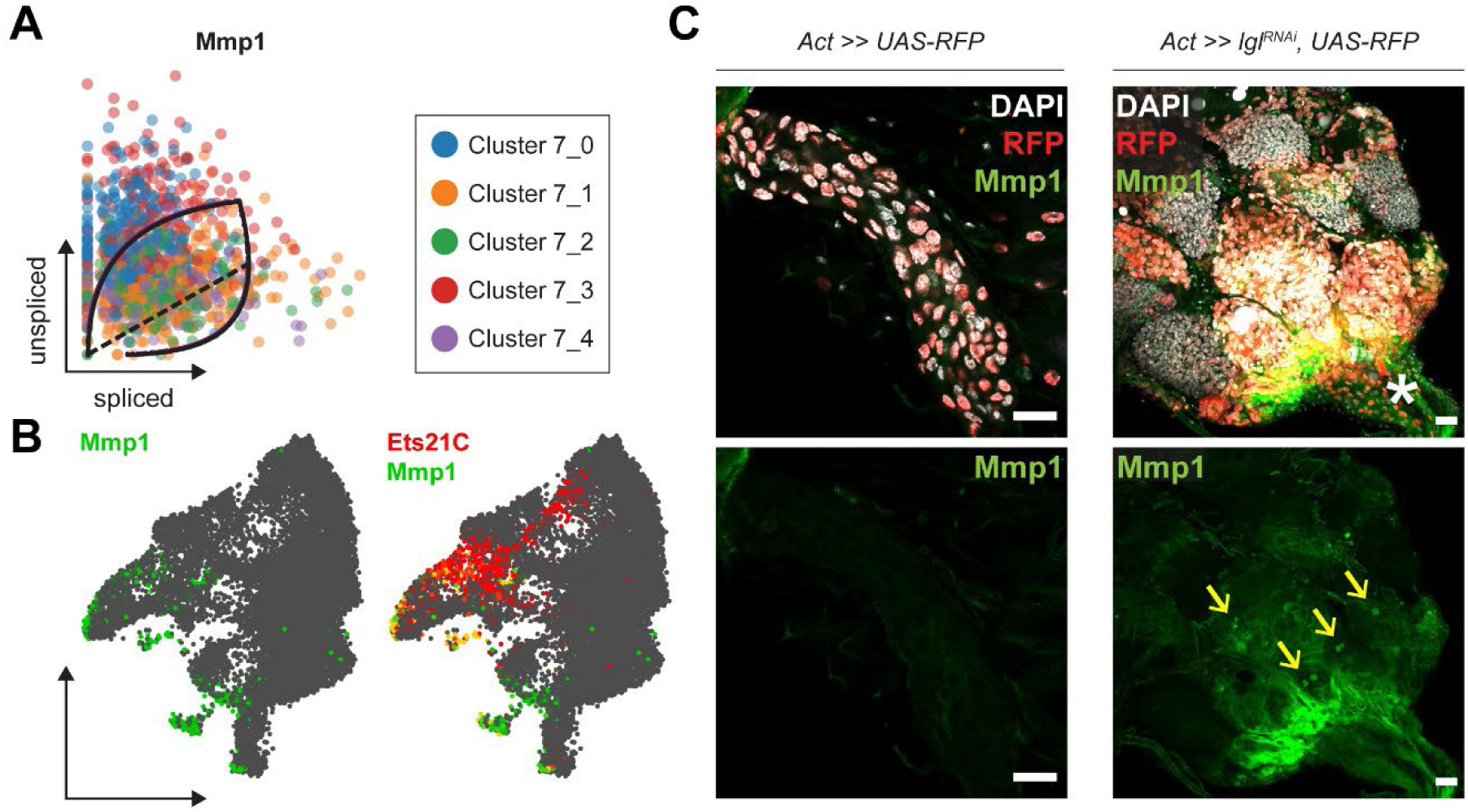
Mmp1 expression in Lgl-KD cells. **A.** Phase portraits of Mmp1 mRNA distribution in cluster 7 cells. **B.** Gene enrichment plot showing the overlap of Mmp1 expression (green) in cluster 7 cells at terminal end-points, that also exhibit Ets21C expression (red). Overlap is shown in yellow. **C.** Confocal Images showing Mmp1 staining in terminal follicle cells of control egg-chambers (in Corpus luteum; left column) and those containing Lgl-KD (3 days) follicle cells (right column) that accumulate near the oviduct marked by an asterisk (*). Transgene expression is marked by RFP expression in red. Externally secreted Mmp1 proteins are highlighted by yellow arrows. Scale bars, 20 µm.

**Supplementary Figure 5:**
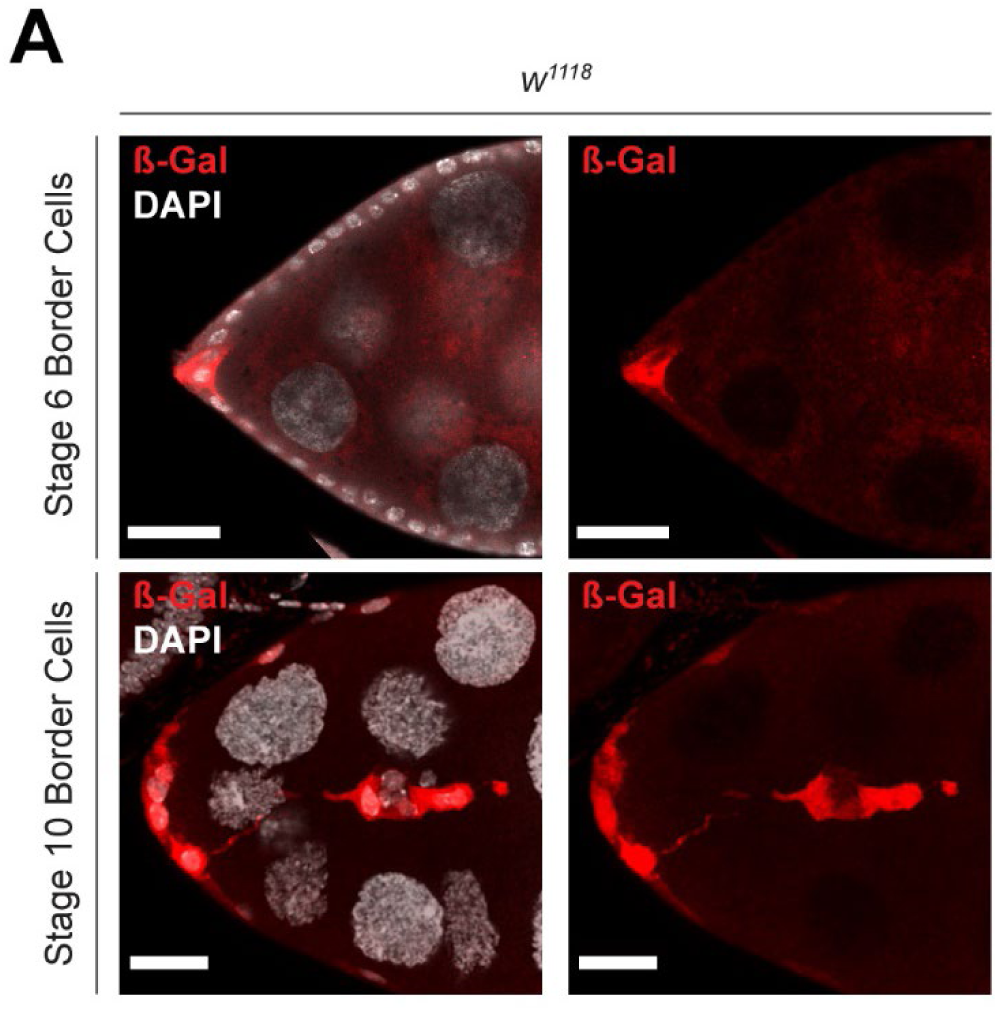
GstD-lacZ expression in *w^1118^* control. **A.** Confocal Images showing GstD-lacZ expression, marked by β-Gal staining (red), in border cells of *w^1118^* egg chambers. Nucleus is marked by DAPI (white). Scale bars, 20 µm.

**Supplementary Figure 6:**
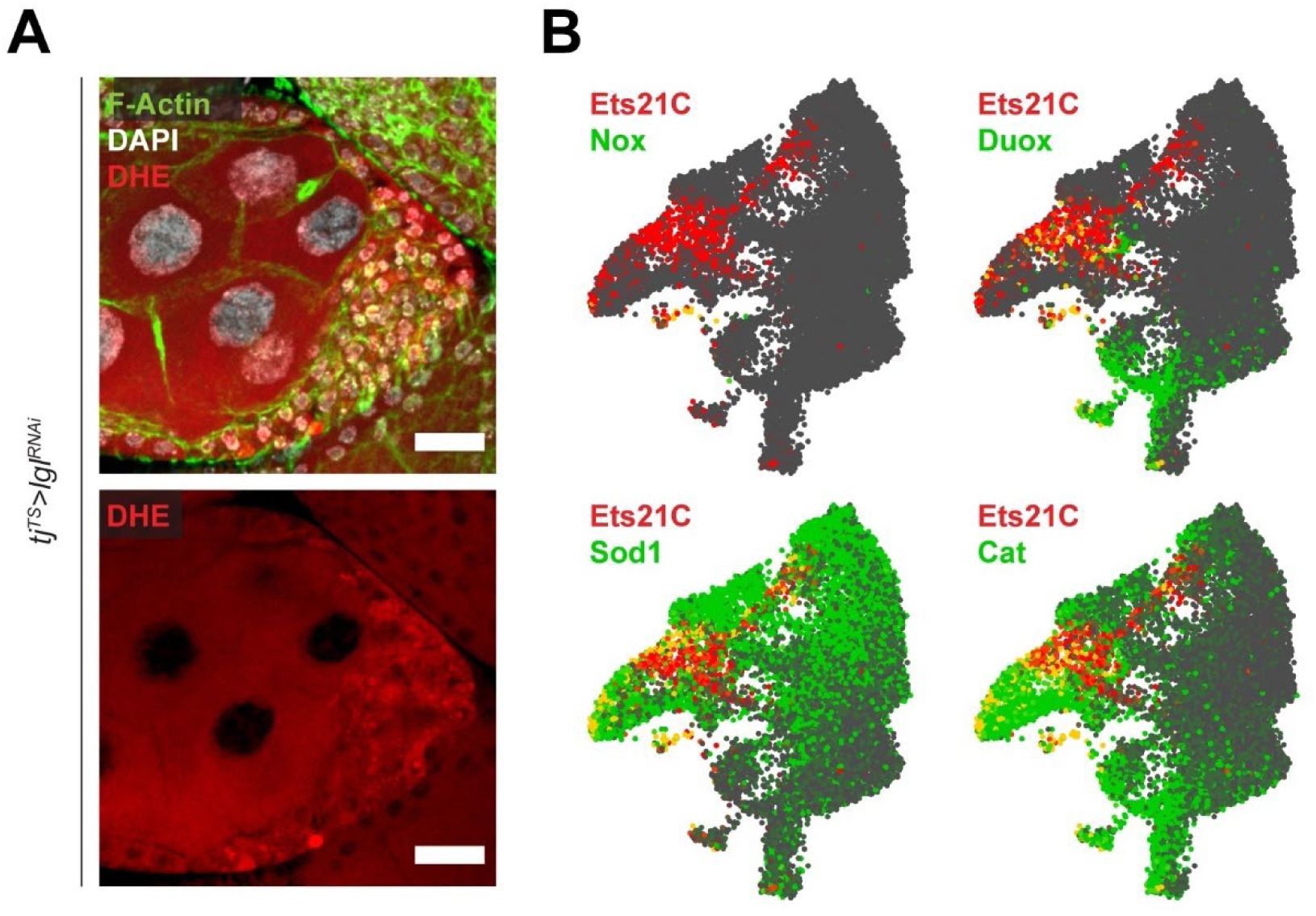
Oxidative-stress response is not detected in cluster 7 cells. **A.** Confocal Images showing dihydroethidium (DHE) staining (red) in the multilayers. Nucleus is marked by DAPI (white). F-Actin is marked by Phalloidin (green). Scale bars, 20 µm. **B.** Gene enrichment plots showing the enrichment overlap of common oxidative-stress response genes (green) with Ets21C (red) that is specific to cluster 7. Overlap of gene expression is observed as cells colored yellow.

## SUPPLEMENTARY DATA LEGENDS

**Supplementary Data 1:** Differentially expressed genes in 96h-Lgl-KD vs Others comparison.

**Supplementary Data 2:** Cluster-specific markers of the *tj^TS^>lgl^RNAi^* (72h) scRNA-Seq dataset.

**Supplementary Data 3:** Regulon Specificity Scores (RSS) of regulons enriched in clusters 7, 8 and 13 and sub-clusters of cluster 7 in *tj^TS^>lgl^RNAi^* (72h) scRNA-Seq dataset.

**Supplementary Data 4:** Differentially expressed genes per subclusters of cluster 7 cells in *tj^TS^>lgl^RNAi^* (72h) scRNA-Seq dataset.

**Supplementary Data 5:** Enriched transcription-factor binding motifs detected in genes expressed in the cells of cluster 7.

**Supplementary Data 6:** Genes showing dynamic behavior of transcription driving the underlying RNA velocity of cluster 7 cells.

